# A new dual-affinity peptide nucleic acid for targeting miRNA-21 precursor rescues tumor repressor PTEN expression

**DOI:** 10.1101/2025.08.08.668835

**Authors:** Yun Lian, Anni Wang, Ruiyu Lin, Xin Ke, Xuan Zhan, Rongguang Lu, Tingli Qian, Arpita Ghosh, Alan Ann Lerk Ong, Desiree-Faye Kaixin Toh, Kiran M. Patil, Manchugondanahalli S. Krishna, Clément Dezanet, Jean-Luc Decout, Souvik Maiti, Hongwei Wang, Gang Chen

## Abstract

MicroRNAs (miRNAs) have multiple functions in cells and are related to many diseases including cancer by regulating posttranscriptional gene expression. The microRNA precursors (pre-miRs), which can be cleaved by Dicer endonuclease to produce mature miRNAs, often have a cleavage site consisting of both double-stranded (ds) and single-stranded (ss) RNA structures. Peptide nucleic acid (PNA) is a kind of analogue of DNA which can hybridize to DNA or RNA through Watson-Crick and Hoogsteen pairing. We previously reported a novel dual-affinity PNA (daPNA) platform that can simultaneously form a duplex and a triplex with a target RNA’s ssRNA-dsRNA junction region. The PNA-RNA complex structure is stabilized by forming antisense PNA (asPNA)-ssRNA duplex immediately adjacent to a chemically modified dsRNA-binding PNA (dbPNA)-dsRNA triplex. In this study, we further explored the application of the daPNA platform to target the precursor of miR-21, which is considered an oncogene. We have designed a set of PNAs including asPNAs, dbPNAs, and daPNAs. The nondenaturing polyacrylamide gel electrophoresis (PAGE) and biolayer interferometry (BLI) data reveal that daPNA-21-10 can strongly bind to pre-miR-21 with high specificity. The data show that pre-miR-21 is easily targeted by traditional antimir, asPNA or dbPNA and optimization of the dsRNA-ssRNA junction position is needed for identifying a tightly binding daPNA. Furthermore, daPNA-21-10 not only inhibits the Dicer activity on pre-miR-21 in cell-free assays, but also down-regulates the expression of miR-21, rescuing PTEN protein expression in cells. Taken together, the biofunction and programmability of daPNA make it a promising platform for probing and regulating miRNA biogenesis and many other RNA-involved biological processes.

**Highlights:**

- Simultaneous recognition of dsRNA and ssRNA regions
- RNA targeting by dividing and conquering through a dsRNA-ssRNA junction construction
- Substrate-specific inhibition of Dicer acting on pre-miR-21
- Derepressing PTEN expression by inhibiting miR-21 maturation

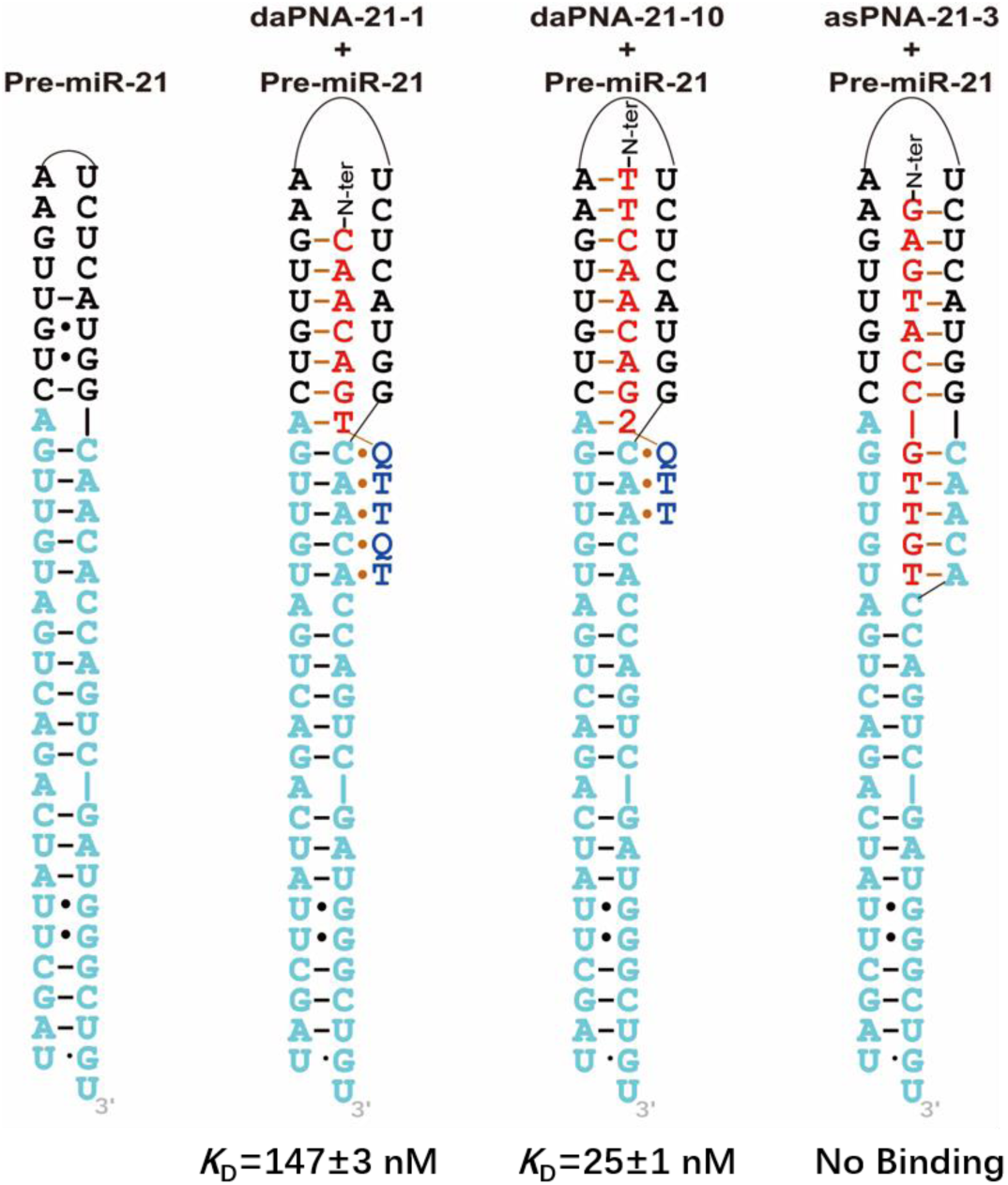

**eToc Blurb:** Lian et al. show that the miR-21 precursor structure containing the Dicer cleavage site can be targeted by a new type of dual-affinity peptide nucleic acids (daPNAs) through the creation of a dsRNA-ssRNA junction, energetically optimized for the simultaneous duplex and triplex formation. Pre-miR-21 structure, which is hardly accessible to traditional antisense strategies and triplex formation alone, can be targeted by a daPNA with high sequence/structure specificity and strong binding affinity. daPNA platform has a great potential in probing many other RNA structures and broad therapeutic applications.

## INTRODUCTION

RNAs play a wide range of roles in catalytic and regulatory activities in cells^1–3^. The functions of RNA are heavily determined by their secondary and tertiary strctures^4–7^. These intricate structures facilitate molecular recognition by small molecules and peptides/proteins^8–10^.

MicroRNAs (miRNAs, **Figure 1**) are a class of small, non-coding RNA molecules that play a crucial role in regulating gene expression at the post-transcriptional level and the dysregulation of miRNAs cause many diseases^11–17^. The maturation process of miRNAs involves multiple steps and enzymatic reactions, transforming primary miRNA transcripts (pri-miRNAs) into precursor miRNAs (pre-miRNAs) as intermediates and mature miRNA molecules that are typically ∼22 nucleotides in length^18,19^. The efficiency and specificity of miRNA maturation are influenced by various factors, including the sequence and structure of the pri- and pre-miRNAs, the expression and activity of the processing enzymes (Drosha, DGCR8, and Dicer), and interactions with regulatory proteins^17,20^. Among all miRNAs, microRNA-21 (miR-21) is well studied as an oncogenic miRNA and is over expressed in many cancers such as non-small cell lung cancer, glioblastoma, and pancreatic adenocarcinoma, etc^21–26^.

**Figure 1.**
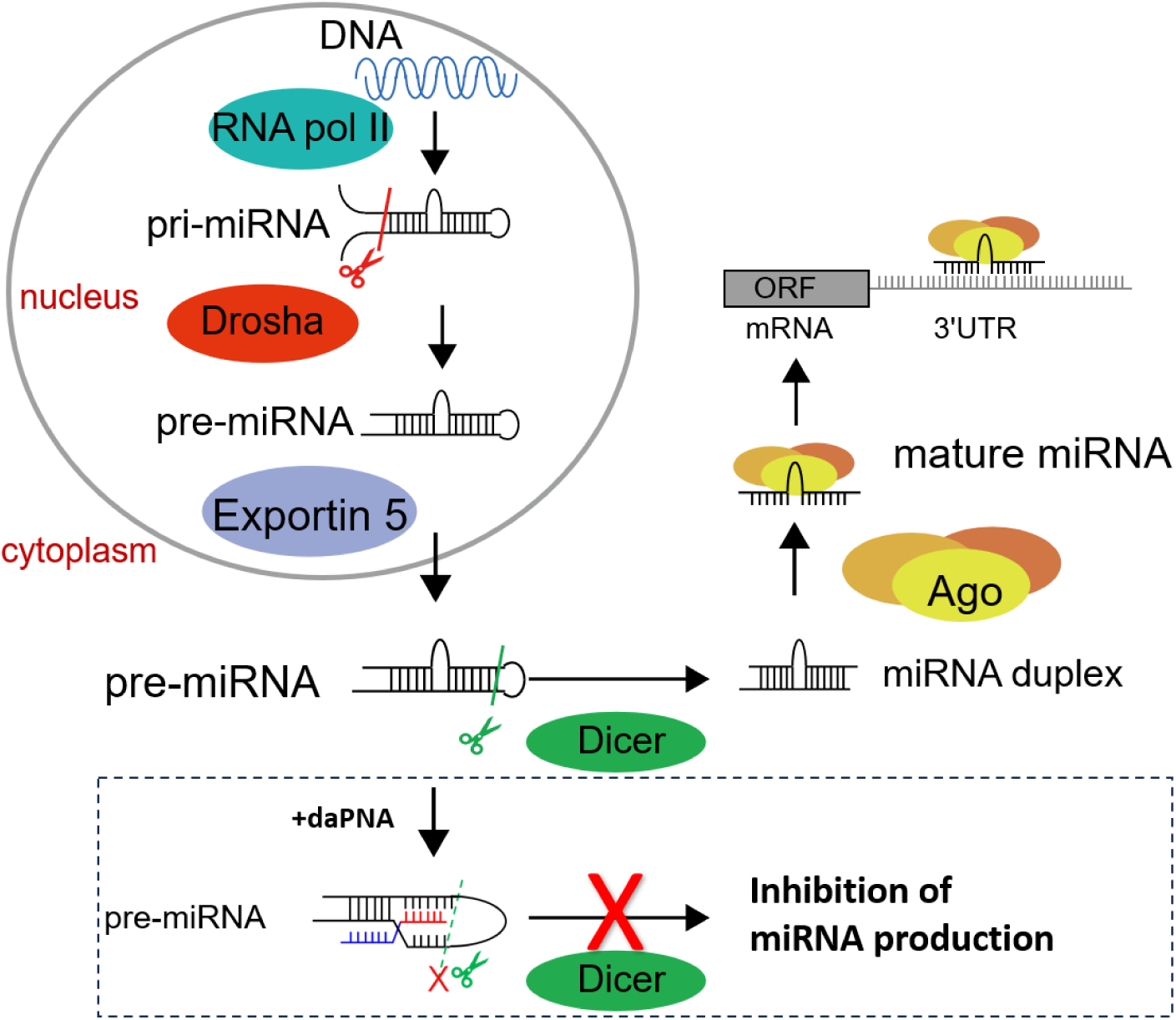
Schematic of the maturation process of miRNA and its inhibition by daPNA targeting pre-miRNA structure through the inhibition of Dicer activity. Binding of miRNA to target mRNA 3’ UTR results in the translation repression. RNAs are shown in black. daPNA is shown in blue and red (see Figure 2, Figure 3). ORF: open reading frame. DGCR8: DiGeorge syndrome critical region 8 Microprocessor Complex Subunit. TRBP: transactivation response element RNA-binding protein. 3’ UTR: 3’ untranslated region.

Antisense oligonucleotides (ASOs), typically ranging from 13 to 25 nucleotides in length, are a major class of RNA-targeting molecules that bind to RNAs including mature miRNAs through Watson-Crick pairing^14,15,27–29^. Some ASOs have been approved for treating different kinds of diseases. Despite their potential, reported applications of programmable structure-specific RNA ligands targeting double-stranded RNAs are relatively limited. Triplex-forming oligonucleotides (TFOs) are one traditional approach that can bind to the major groove of RNA duplexes (double-stranded RNAs, dsRNAs) and form triplexes through Hoogsteen pairing^30–32^. However, due to charge repulsion among the negatively charged backbones in a triplex, TFOs often have relatively weak binding affinity comparing to ASOs. The difficulty in targeting inverted base pairs such as C-G and U-A through natural nucleobases makes the application of TFOs even more limited^33,34^. In addition, TFOs often exhibit stronger binding to double-stranded DNA (dsDNA) than to dsRNA. To enhance the properties of TFOs binding to dsRNAs, modifications have been made to their sugar-phosphate backbones and nucleobases^35^.

Among nucleic acid analogues, peptide nucleic acid (PNA) is distinguished by its pseudopeptide backbone (**Figures 2**, **3**), which provides significant chemical stability and resistance to nucleases^36–40^. Neutral PNA backbone also gives PNA-DNA/RNA duplexes (**Figure 2a**) higher stability than DNA-DNA or RNA-RNA duplexes^41–43^. Notably, there are a variety of artificial nucleic acids including PNAs that have demonstrated bioactivity in animal models and have promising roles as biosensors and therapeutics^37,44–47^. It’s worth noting that since natural dsRNAs are usually not fully complementary, which allows antisense PNAs (asPNAs) to invade the relatively weakly formed dsRNA region and form asPNA-RNA duplexes (**Figure 2a**).

**Figure 2.**
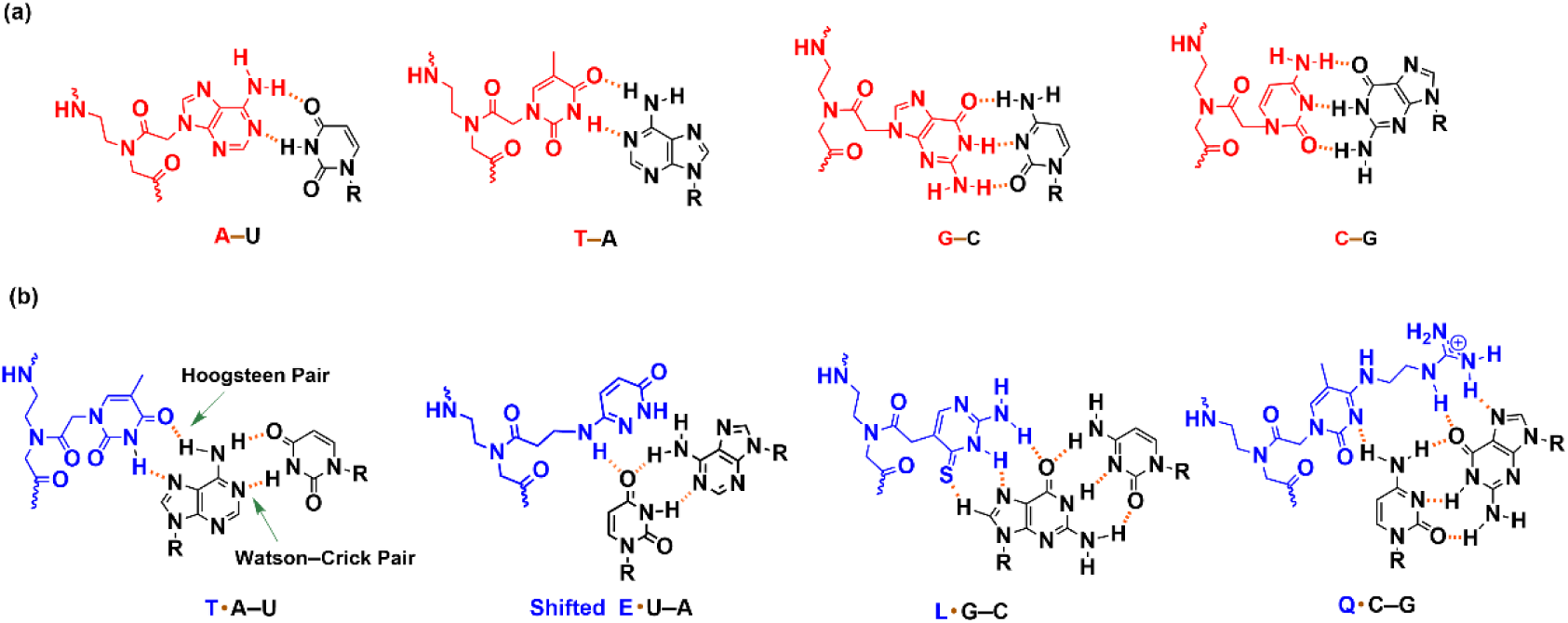
Chemical structures of base pairs and base triples formed between PNA and RNA. (a) PNA-RNA base pairs of A-U, T-A, G-C and C-G formed by asPNA and ssRNA. (b) dbPNA•dsRNA base triples of T•A-U, Shifted E•U-A, L•G-C, Q•C-G^60^ formed by dbPNA and dsRNA. L monomer shows enhanced recognition of G-C pair compared to C monomer with reduced pH dependence^68^. The shifted E•U-A triple is drawn in a geometry^60^ based on the previous study on backbone-base linker optimization^69^. The Watson-Crick pairs are shown as orange lines and Hoogsteen pairs as orange dots. dbPNAs are shown in blue and asPNAs in red. RNAs are shown in black.

**Figure 3.**
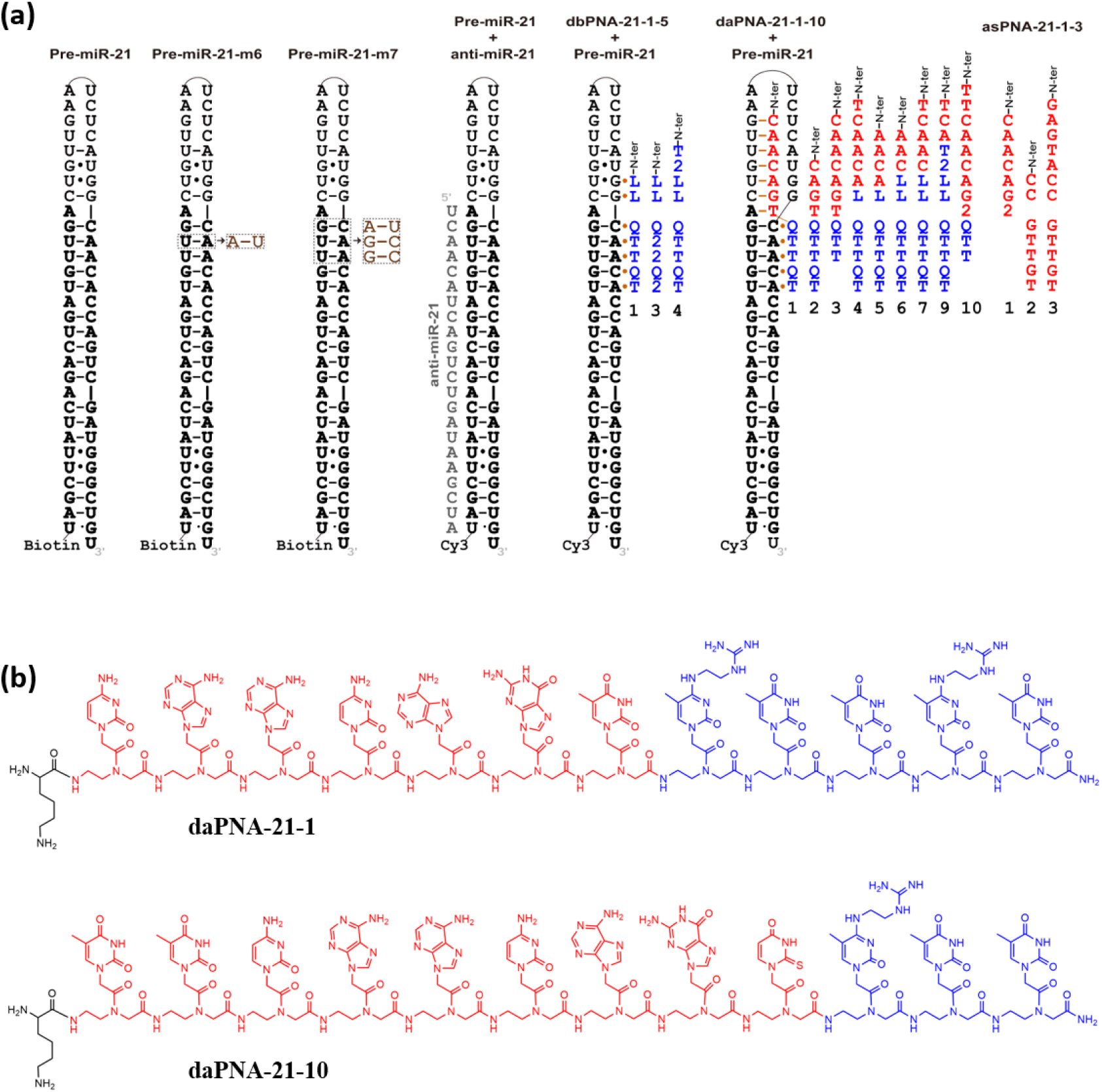
Sequences and structures of PNAs for binding pre-miR-21. (a) RNA constructs and potential binding by PNAs and a 2’-OMe modified anti-miR-21. The Watson-Crick pairs are shown as lines and Hoogsteen pairs as orange dots. dbPNAs are shown in blue and asPNAs red. RNAs are shown in black, with the mature miR-21 sequences shown in bold. Only daPNA-21-1 and daPNA-21-10 show strong binding. (b) Chemical structures of daPNA-21-1 and daPNA-21-10. The dbPNA and asPNA monomers are shown in blue and red, respectively. The N-terminal lysine residue is in L configuration.

We and others have previously shown that some modified PNAs with modified nucleobases (dbPNA, short for dsRNA-binding PNA, **Figure 2b**) can form dbPNA·dsRNA triplex structures with dsRNA^35,48–60^. dbPNAs show high binding specificity to dsRNA with reduced binding to dsDNAs and ssRNAs. Though dbPNAs have shown biological activity in various cellular processes including translation, miRNA maturation and RNA editing^60^, the effectiveness and availability of dbPNAs are limited because most natural functional dsRNAs are not fully complementary, but often interrupted by ssRNAs. In fact, targeting to an RNA region often requires a simultaneous binding to both dsRNA and ssRNA.

Based on the preferential binding of asPNAs and dbPNAs to ssRNA and dsRNA, respectively, we have previously developed a novel dual-affinity PNA (daPNA) platform which includes both asPNA and dbPNA, aiming to target RNA junction regions composed of both ssRNA and dsRNA ^18^. In this study, we further developed daPNAs for targeting pre-miR-21 structure (**Figure 3**). Binding studies indicate that the optimized daPNAs can effectively target dsRNA-ssRNA junctions in pre-miR-21, and exhibit biological activities in both cell-free and cell culture assays.

## RESULTS AND DISCUSSION

### Binding properties of daPNAs targeting pre-miR-21 with varied dsRNA-ssRNA junction positions

Based on the nuclear magnetic resonance (NMR) structure of pre-miR-21^23^, there are potentially at least four dsRNA-ssRNA junctions that may be utilized for daPNA binding (see **Figure 3a, S1a**). Based on the sequence and structure of pre-miR-21, we designed a series of dbPNAs, asPNAs, and daPNAs composed of asPNA part forming a fully complementary duplex with ssRNA in pre-miR-21, and dbPNA forming a parallel triplex with dsRNA (**Figure 3, S1a**). Varied PNAs were made (**Figure S1a,S2**) to study the best binding mode with a junction between ssRNA and dsRNA optimized for the PNA-RNA duplex/triplex complex structure formation. The PNAs were designed to form base pairs or base triples with the RNA (**Figure 2 and Figure S1b**). High performance liquid chromatography (HPLC) and Mass Spectrometry (MS) were used to characterize the purity and identity of the PNAs (**Figure S2**).

To study the binding affinity of PNAs towards pre-miR-21, we conducted non-denaturing polyacrylamide gel electrophoresis (PAGE) (**Figure 4a,S3**)^61^. The results suggest that daPNA-21-1 (*K*_D_=270 nM) and daPNA-21-10 (*K*_D_=26 nM) can efficiently bind to pre-miR-21 (**Figure 4a, c, S3**). daPNA-21-3, daPNA-21-4 and daPNA-21-7 show relatively weak or comparable binding affinities compared to daPNA-21-1. To further characterize the binding properties, we conducted bio-layer interferometry (BLI) (**Figure 4b,c, S4**)^62,63^. The BLI data are generally consistent with those of PAGE with comparable equilibrium dissociation constants (**Figure 4c, Table S1**). The BLI data reveal that daPNA-21-10 has an improved binding affinity, mainly through an increase in on rate, indicating that lengthening the asPNA segment may facilitate the initial binding of the daPNAs.

**Figure 4.**
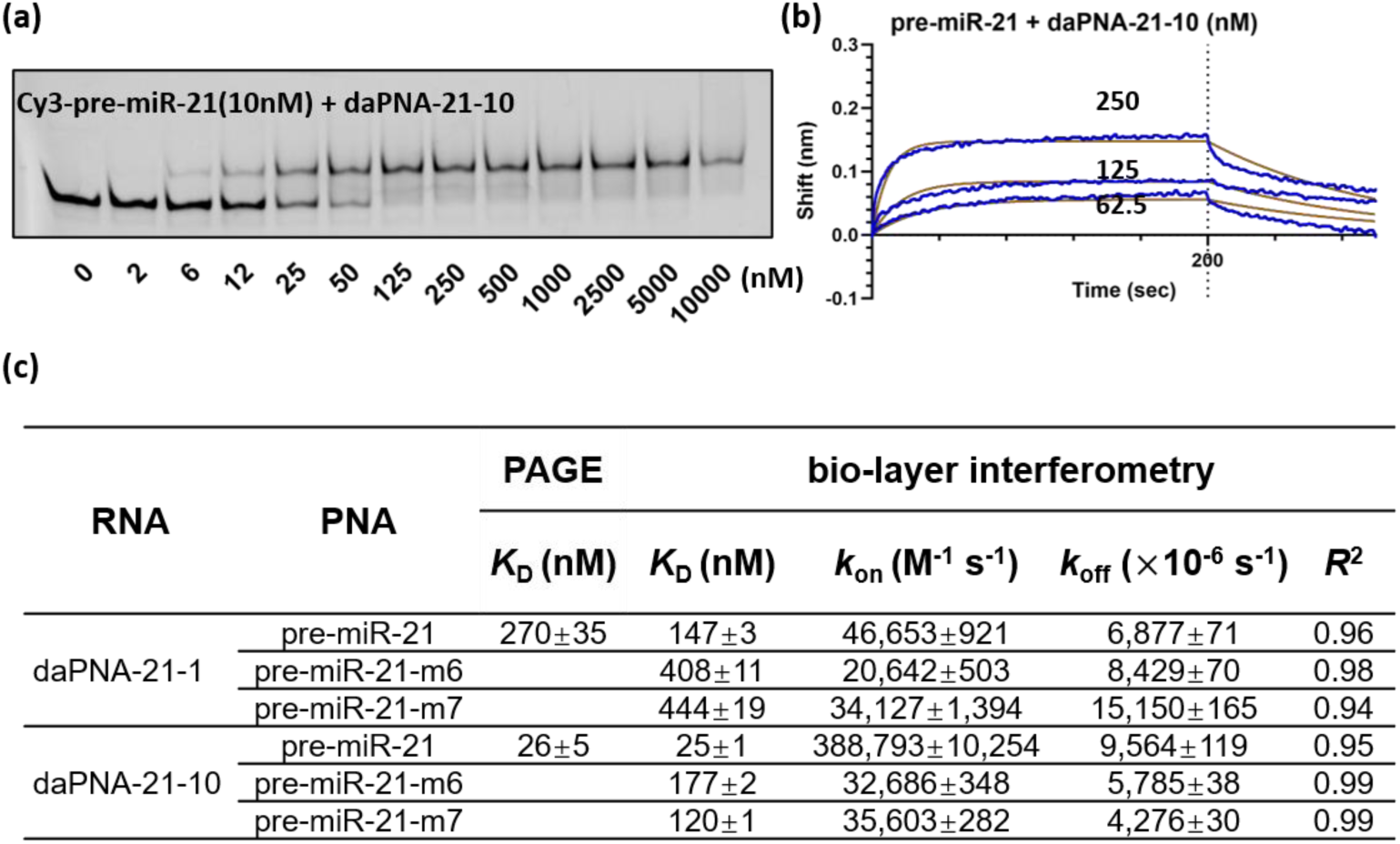
Targeting pre-miR-21 with daPNAs. (a) Nondenaturing PAGE binding assay data for daPNA-21-10 targeting Cy3-pre-miR-21. (b) BLI data for daPNA-21-10 targeting biotin-pre-miR-21. (c) Summary of BLI and PAGE data for daPNA-21-1, 10 targeting pre-miR-21. NB: no binding observed. More binding data can be found in Figure S4 and Table S1. The other PNAs shown in Figure 3a typically show much weakened or no binding.

Besides, we tested the binding affinities of asPNAs (**Figure 3c, S4,5**) to prove the necessity of dbPNA presence in daPNA. As shown in PAGE, asPNA-21-1, asPNA-21-2, and asPNA-21-3 showed weak or no binding to Cy3-pre-miR-21 (**Figure S5**). This suggests that the existence of dbPNA part in daPNA is crucial for targeting RNAs. It is likely that the dbPNA-binding dsRNA region is highly stable and not easily invaded by asPNAs.

We further constructed a series of mutants of pre-miR-21 to test if modulating the structure and stability above the A bulge affects the daPNA binding (**Figure S6**). Pre-miR-21-m1 has a G-U pair replaced by a G-C pair, which is expected to result in the stabilization of the structure and inhibit the asPNA binding. Indeed, the PAGE data show no binding of the tested daPNAs to pre-miR-21-m1. We observed no improved binding by daPNA-21-6, daPNA-21-7, and daPNA-21-9, even though they are expected to form an L•G-C base triple with the mutated G-C pair. It is probable that the G-U wobble pair to G-C Watson-Crick mutation stabilizes the nearby region above the mutated base pair, resulting in the inhibition of binding by the asPNA segments of daPNA-21-6, daPNA-21-7, and daPNA-21-9. Similarly, the mutant pre-miR-21-m4 (with the deletion of the A bulge) also shows no binding, presumably because the single residue deletion results in a loss of asPNA-RNA base pair as well as stabilizing the nearby RNA structure preventing the invasion by asPNAs. As expected, a destabilizing mutation in pre-miR-21-m2 (G-U to C•U mismatch) and pre-miR-21-m3 (G-C to A•C mismatch) allows the binding by daPNA-21-1, daPNA-21-3, and daPNA-21-4, presumably because the structure-destabilizing mutations facilitates the invasion by the asPNA segments. The data show that daPNAs may be used to probe local structural stabilities and dynamics in RNAs.

We also conducted a PAGE assay studying the influence of temperature on daPNA-21-1 and a 2’-OMe modified anti-miR-21 (**Figure 3a**) binding to RNAs (**Figure S7**). It turned out that while the binding of anti-miR-21 depends heavily on the incubation temperature, daPNA-21-1 shows a stable binding to Cy3-pre-miR-21. This feature is probably caused by dbPNA part in daPNA, since it does not require a high temperature to break the Watson-Crick pairs in the pre-miR-21 to allow the access for anti-miR-21^55^. The data clearly show that daPNA is advantageous in targeting structured RNAs, with the dbPNA and asPNA segments recognizing locally rigid and flexible regions of RNA, respectively.

Our data suggest that the A bulge position may be ideal for generating the dsRNA-ssRNA junction for daPNA binding. For example, with the A bulge residue recognized an asPNA residue T, as a replacement of asPNA-RNA T-A pair with a dbPNA-dsRNA L•G-C base triple in the junction point causes a weakened binding **(Figure 4c, S3,S6, Table S1**). The A bulge is on the nonpurine-rich strand (opposite to the middle strand), which makes it difficult to be recognized by a dbPNA residue. It was previously reported that an A bulge on the purine-rich strand (middle strand) may be recognized by a T base within a dbPNA through the Hoogsteen T•A pair formation^64^. We further tested if dbPNA•RNA-RNA triplex is formed or not in the binding process by designingpre-miR-21-m6 and pre-miR-21-m7, with one A-U pair and all three Watson-Crick pairs recognized by the dbPNA segment of daPNA-21-10 mutated (**Figure 3a**). BLI data show that pre-miR-21-m6 and pre-miR-21-m7 show significantly weakened binding for both daPNA-21-1 and daPNA-21-10 (**Figure 4c,S4, Table S1**). Interestingly, the BLI data suggest that, compared that of daPNA-21-1, the on rate of daPNA-21-10 is significantly reduced upon the mutation from pre-miR-21 to pre-miR-21-m6 and pre-miR-21-m7, which suggests that the a matching dbPNA segment is critical for initiating the binding of daPNAs to pre-miR-21.

### daPNA-21-10 inhibits Dicer cleavage *in vitro*

To test if daPNAs can inhibit miRNA maturation, we conducted Dicer cleavage assay. The results show that daPNA-21-10 can specifically inhibit Dicer cleavage on pre-miR-21 *in vitro* (**Figure 5**). Meanwhile, daPNA-21-4, 7, 9 and dbPNA-21-1 showed no obvious inhibition in Dicer cleavage (**Figure S8**). This is consistent with the weak binding shown in PAGE, which indicated that Dicer cleavage efficiency highly depends on the binding affinity. It is worth to be noted that daPNA-21-1showed aggregation with RNA^65^ in both PAGE Dicer cleavage assay (**Figure S8**) and PAGE binding assay (**Figure S3**) at concentration above 5 μM. Taken together, the binding and cell-free Dicing data show that daPNA-21-10 may be a top candidate sequence for further studies.

**Figure 5.**
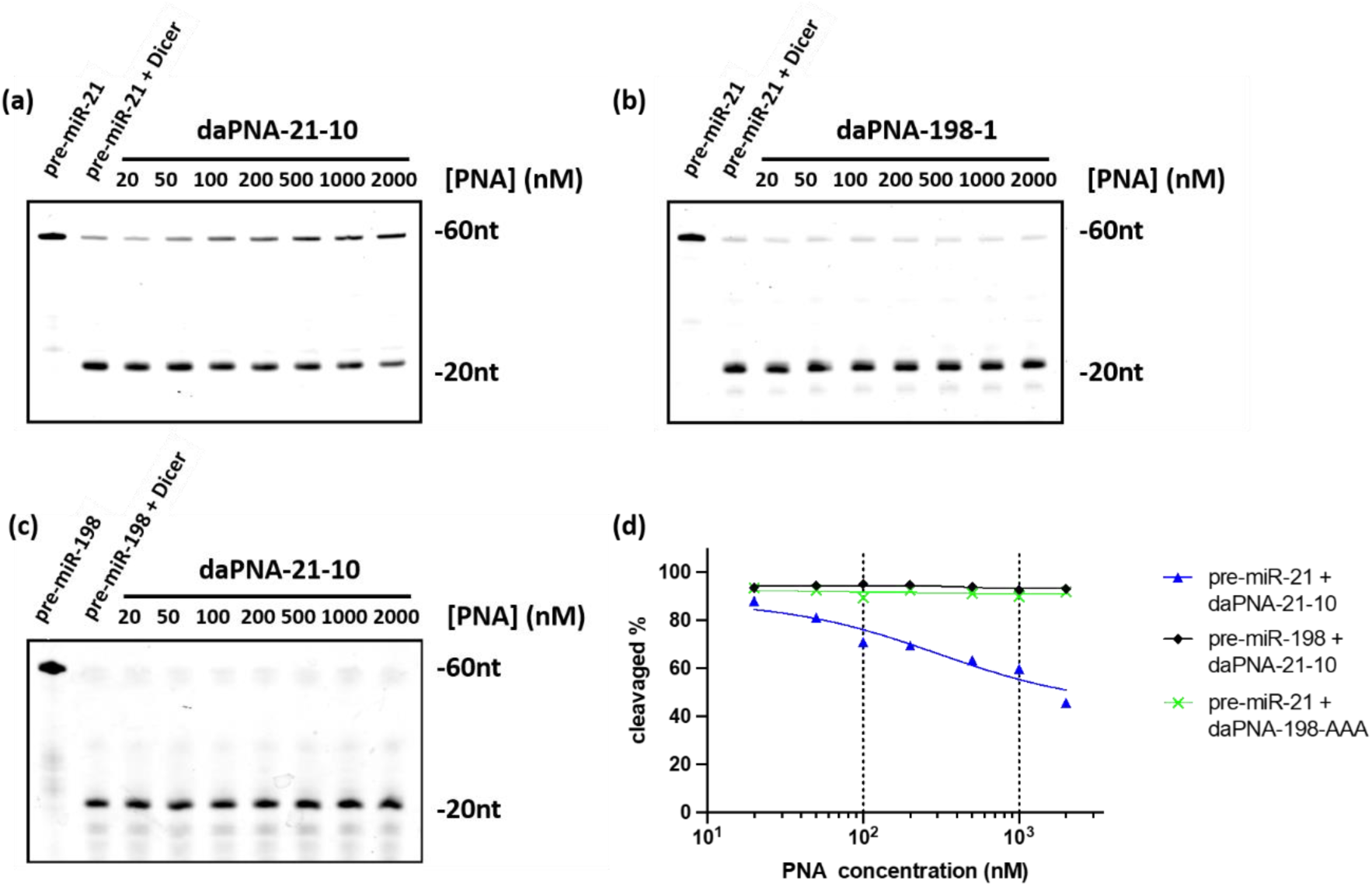
Dicing assay for daPNA-21-10 and pre-miR-21-HP. (a-c) Dicer cleavage assay data for daPNA-21-10 inhibition of Dicer cleavage of pre-miR-21. daPNA-198-1 and pre-miR-198 were used as control PNA and RNA, respectively. Gradient PNAs were pre-incubated with RNA at 50℃ for 10 min, then slow cooling to RT. After adding hDicer, the dicing system was incubated at 37℃ for 90 min. The reaction was stopped by boiling with 2× RNA loading buffer containing 8 M Urea, 1× TBE, 0.05% bromophenol blue for 10 min followed by ice bathing. RNA products were then analyzed on 15% polyacrylamide, 8 M urea denaturing gel electrophoresis. (d) Summary of the Dicer cleavage data.

### daPNA-21-10 inhibits miRNA-21 expression in cell cultures

To test the bioactivity of daPNAs in the cell, we conducted a dual-luciferase reporter assay in HEK293T cell (**Figure S9a**). The results suggest that daPNA-21-1 at 0.5μM can derepress the inhibition effect of miR-21 on expression of firefly luciferase (**Figure S9b**). We applied daPNA-21-10 in HEK293T and Hela cells to further confirm the effect of substrate-specific inhibition of Dicer activity on pre-miR-21. The cytotoxicity was determined through CCK8 assay and daPNA-21-10 showed no obvious cytotoxicity as high as 50 μM in HEK293T cells (**Figure 6a**). We conducted qPCR to detect the expression level change of related RNAs including pri-miR-21, pre-miR-21, miR-21 and PTEN mRNA in HEK293T and Hela cells treated with daPNA-21-10. The data reveal that daPNA-21-10 can indeed inhibit Dicer cleavage in both HEK293T and HeLa cels, resulting in an increase and decrease of the levels of pre-miR-21 and miR-21, respectively(**Figure 6b, c**), while the control daPNA-198 shows no effect on the expression levels of the RNAs (**Figure 6d**). Interestingly, pri-miR-21 expression does not change significantly in HeLa cells (**Figure 6c**), suggesting that the activity of Drosha is not influenced by daPNA-21-10 application. Besides, PTEN mRNA expression level is insensitive to the application of daPNA-21-10, consistent with the fact that miR-21 binds to the 3’-UTR of PTEN mRNA with partial complementarity^66^ (**Figure S9b**).

**Figure 6.**
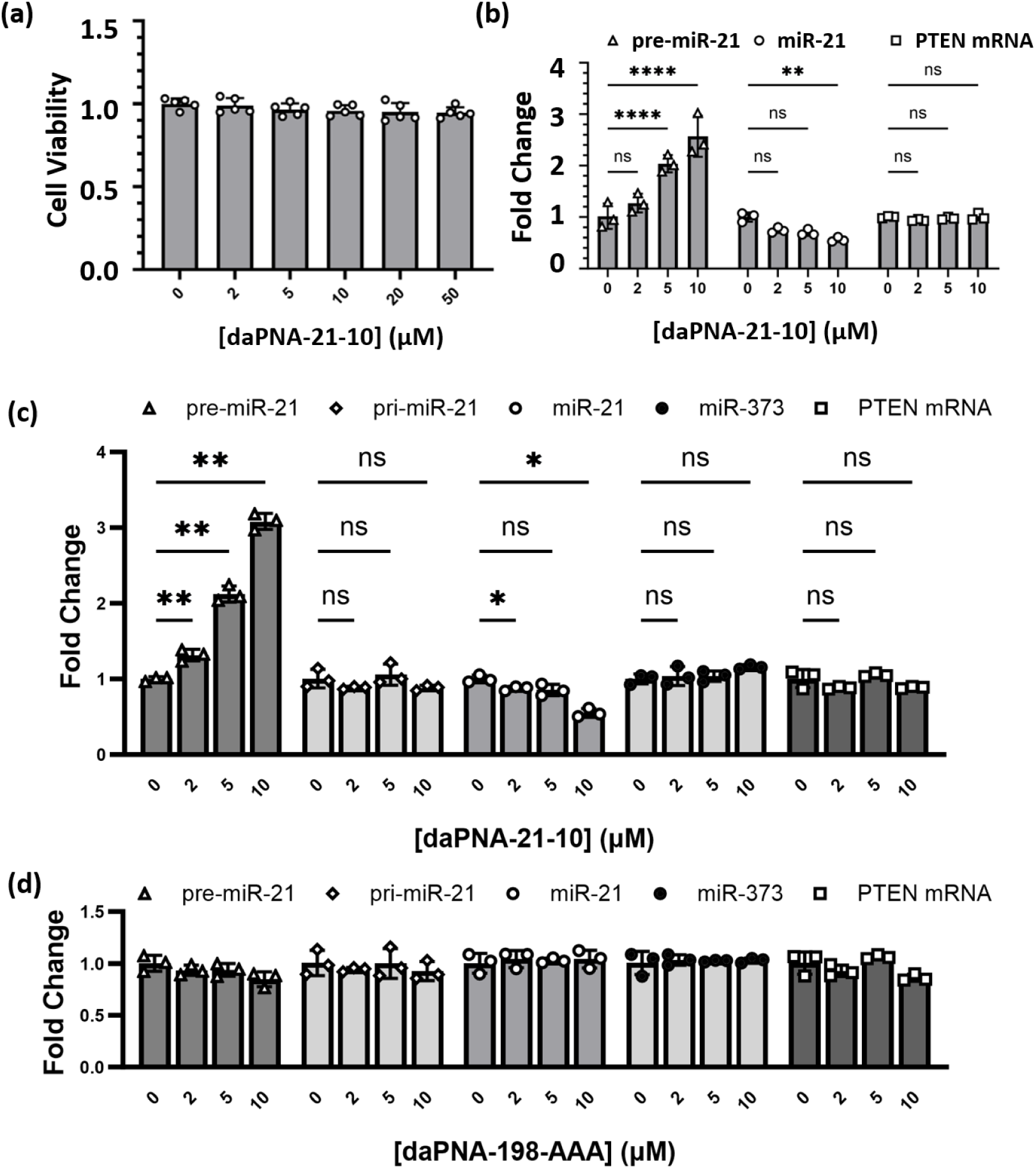
Testing the effects of daPNA-21-10 on cell viability and RNA expression in HEK293T and Hela cells. (a) Cell viability of daPNA-21-10 determined by CCK8 in HEK293T. HEK293T cells were transfected with 0, 2, 5, 10, 20 and 50 μM daPNA-21-10 and treated for 24 h in 96-well plate. (b) Effects of daPNA-21-10 on RNA expression of pre-miR-21, miR-21 and PTEN mRNA in HEK-293T. (c,d) Effects of daPNA-21-10 (c) and control daPNA-198-AAA (d) on RNA expression of pre-miR-21, pri-miR-21, miR-21, miR-373 and PTEN mRNA in HeLa. The cells were transfected with different concentrations of daPNA-21-10 and treated for 72 h in 24-well plate. The variance is analyzed using Two-Way ANOVA. The error bars represent ± S.D. * P < 0.05, ** P < 0.01, *** P < 0.001, **** P < 0.0001, ns: not significant.

### daPNA-21-10 rescues PTEN protein expression in cells

According to the results of real-time qPCR, the miRNA level can be downregulated to about 50% the normal level if daPNA-21-10 was used at 10 μM and treated for 72 h. We applied the same conditions here for the Western Blot assay to study the effect of daPNA-21-10 on PTEN protein expression level. The data suggest that application of daPNA-21-10 results in an increased PTEN expression by ∼70% compared to the vehicle group (**Figure 7**). Antimir-21 and miR-21 mimic showed enhancing and inhibition effect on PTEN protein expression, respectively, as expected. The data are consistent with the result of real-time qPCR and indicate that daPNA-21-10 can successfully derepress the effect of miR-21 on PTEN expression in HEK293T cells and HeLa cells. We observed relatively higher efficiency of daPNA-21-10 in the inhibition of mature miR-21 in HEK293 than in HeLa, probably because the endogenous miR-21 level is higher in HeLa than in HEK293T^67^. The mature miRNAs are bound to and protected by AGO protein with a relatively long lifetime, thus requiring a relatively long time (72h) to observe the inhibitory effect of miRNA expression.

**Figure 7.**
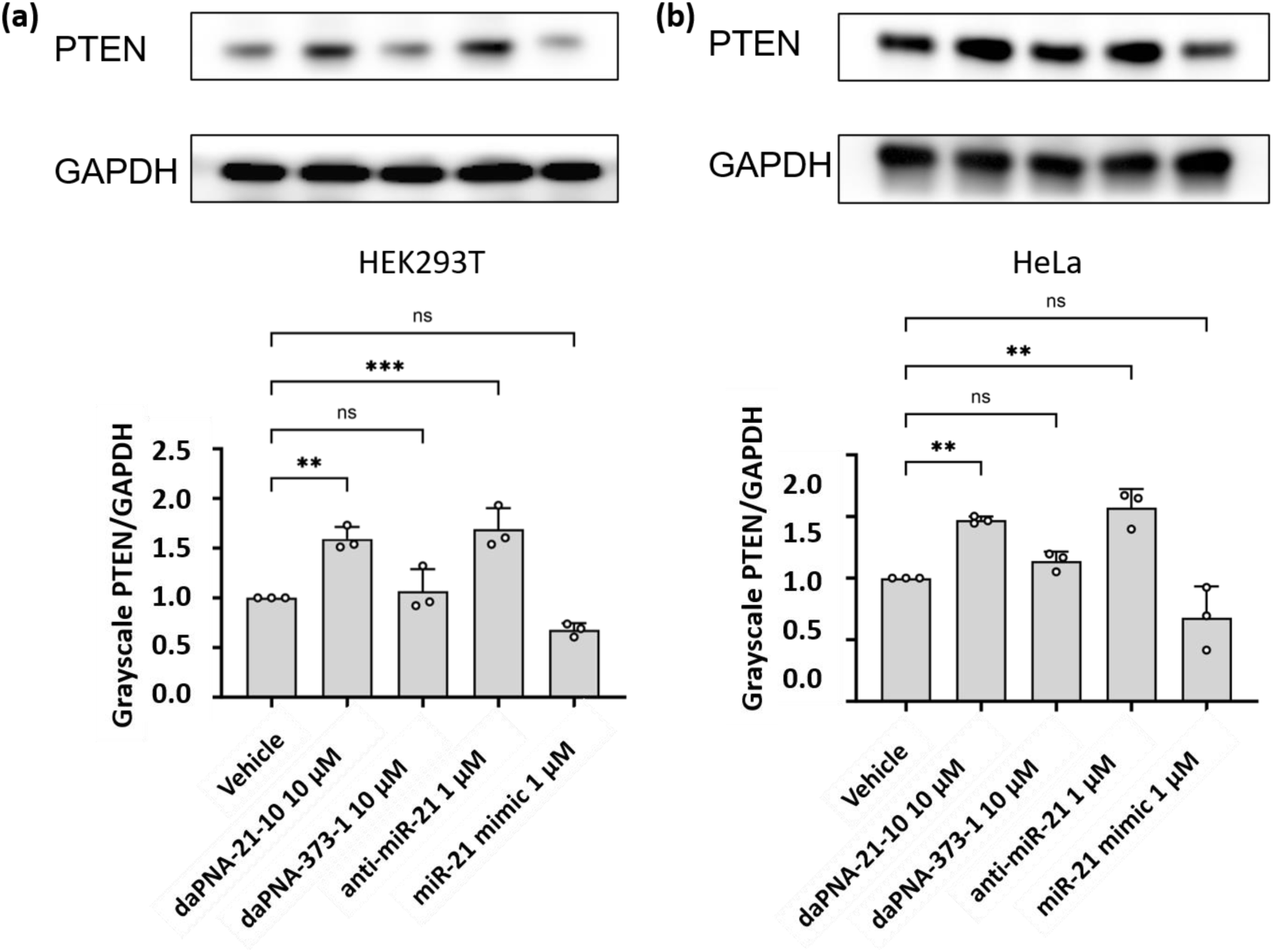
Effects of daPNA-21-10 application on PTEN protein expression in HeLa cells (72h post PNA application). (a) PTEN protein expression in HEK293T cells treated by PNAs. (b) PTEN protein expression in HeLa cells treated by PNAs. Cells were seeded in 24-well plate at 50,000/well. PNAs or anti-miR/mimics were transfected to a final concentration as indicated for 72 h. The data were analyzed by GraphPad Prism 9.3 and calculated by an ordinary one-way analysis of variance (ANOVA) using Dunnett’s multiple comparisons test against the mean of plasmid alone group. The error bars represent ± S.D. * P < 0.05, ** P < 0.01, *** P < 0.001, **** P < 0.0001, ns: not significant.

It is worth noting that there is no obvious cytotoxicity in HEK293T (50 μM, **Figure 6a**) and HeLa (10 μM) cells upon the treatment with daPNA-21-10, although PTEN is a repressor of cell proliferation. In fact, miR-21 has different effects on different cells including many cancer cell lines. Further work is needed to test the dose response of daPNA-21-10. In addition, we expect that the intracellular uptake of the PNAs may need to be improved. Our proof-of-concept work reported here clearly provides the foundation for future work on further developing daPNAs as useful regulators of miRNA biogenesis and functions.

### Conclusions

In summary, we discovered that the construction of daPNA by the combination of asPNA and dbPNA can successfully facilitate the high-specificity and high-affinity targeting to the dsRNA-ssRNA junction region in miRNA-21 precursor with both duplex and triplex formed simultaneously. The binding of daPNA-21-10 to pre-miR-21 results in the substrate-specific inhibition of Dicer cleavage and miRNA-21 maturation both *in vitro* and in cell cultures, and recovers the target protein PTEN expression. The new binding mode proved to be programmable and efficient in targeting complex RNA structures where both ssRNA and dsRNA are present, although the dsRNA-ssRNA junction position may require optimization through the adjustment of the dbPNA-asPNA segments in a daPNA. Successful design of a daPNA may benefit from a deep understanding of the energetics of dbPNA segment binding without disrupting pre-existing dsRNA structure and the disruption of the adjacent RNA structures to allow the formation of asPNA-ssRNA duplex structure. The daPNA platform has the potential as a general therapeutic targeting strategy for other miRNA-related diseases, and as dsRNA-ssRNA junction-specific probes to study functions of various noncoding RNAs with complex structures.

## EXPERIMENTAL PROCEDURES

### Synthesis of PNAs and RNA oligomers

The RNAs labelled with Cy3 or biotin were purchased from Biosyntech China and QingKe Bio (China), respectively. PNA monomers including Adenine (A), Guanine (G), cytosine (C) and thymine (T) were purchased from QYAOBIO, China. PNA monomers L and Q were synthesized per the methods reported^49,68^. PNA oligomers were synthesized manually via Solid-Phase Peptide Synthesis (SPPS) method. The PNA oligomerization was tested by Kaiser test. PNA oligomer cleavage was done by trifluoroacetic acid (TFA) and trifluoromethanesulfonic acid (TFMSA) method. The oligomers were then precipitated with diethyl ether, dissolved in deionized water and purified with water-CH3CN-0.1% TFA in reverse-phase HPLC. Characterization is done by LC-MS/MS (Q Exactive HF-X, Thermo Fisher). The concentration of PNAs and RNAs were determined by absorbance at 260 nm obtained by Shimadzu 2550 spectrometer at 65℃/85℃, respectively, and calculated by Lambert-Beer law with extinction coefficients of A, G, C, T, L and Q as 13.7, 11.7, 7.3, 8.8, 7.3 and 7.3 mM^−1^cm^−1^ for PNA monomers, and A, G, C, U as 15.4, 11.5, 7.2 and 9.9 mM^−1^cm^−1^ for RNA bases.

### Nondenaturing polyacrylamide gel electrophoresis

The nondenaturing PAGE binding assay was performed in 20% gel with arc:bis= 37.5:1. The concentration for RNAs used is 10 nM or as indicated. The RNAs attached by fluorophores were diluted in the incubation buffer containing 200 mM NaCl, 0.5 mM EDTA, 20 mM HEPES, pH 7.5, heated at 95℃ for 5 min, and then snap-cooled on ice for 5 min. Then the RNAs were annealed with PNAs at different concentrations by slow cooling from 65℃ to room temperature, or incubated at 37℃ for 2 h if indicated. The samples were then incubated at 4℃ overnight. Before loaded, the samples were added with 35% glycerol (20% of the total volume) followed by vortex. The gels were then run at 4℃ for 5 h under 250V in 1× TBE, and then imaged using an Amersham Imager 680.

### Bio-layer interferometry assay

The RNAs labelled with 5′ biotin was diluted to 200 nM in binding buffer containing 200mM NaCl, 0.5mM EDTA, 20mM HEPES, pH 7.5. The PNAs were diluted in binding buffer at various concentrations. The time periods set for binding-loading-binding-association-dissociation are 120, 300, 120, 600 and 600 seconds, respectively. The data were then fitted using the Gator Prime software or GraphPad with association-then-dissociation model by least squares regression and confidence intervals (CI) set as 95%. The formula for association phase and dissociation phase are as follows:

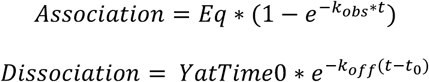

Where *Kobs* = [PNA] ∗ *k*on + *k*off, Eq = Bmax ∗ [PNA]/([PNA] + *K*D) and 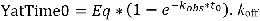 is the dissociation constant in min-1, *k*_on_ is the association constant in inverse minutes multiplied by inverse concentration and *K*_D_ is computed from *k*_off_/*k*_on_ expressed in Molar units. A global fitting is applied and nonspecific binding (NS) is set to be 0.

### *In vitro* dicing assay

Gradient PNAs were pre-incubated with Cy3 labelled pre-miR-21 RNA oligomers at 50℃ for 10 min, then slow cooling to RT. After hDicer was added, the dicing system was incubated at 37 ℃ for 90 min. The reaction was stopped with 2× RNA loading buffer containing 8 M Urea, 1× TBE, 0.05% Bromophenol blue, boiled for 10 min, and subsequently chilled on ice. RNA products were analyzed on 8 M urea denaturing 15% polyacrylamide gel electrophoresis and visualized with a Typhoon Trio Imager (Amersham Biosciences).

### Dual-luciferase reporter assay

HEK-293T cells were cultured in Dulbecco’s modified Eagle’s medium (Gibco) supplemented with 10% FBS (BI), 100 U/ml of penicillin and 100 μg/ml streptomycin (Sangon) at 37℃, 5% CO_2_. The cells were seeded in 96-well plate at 10,000/well. After 24 h, reporter plasmid pmirGLO-miR-21 was transfected to a final concentration of 1 ng/μL, together with pre-miR-21 at 10 nM and different concentrations of PNAs as indicated by Lipo3000 (Invitrogen) per the manual. After 48h, the Firefly and *Renilla* signals were measured by Envision microplate reader (PerkinElmer). The experiment was performed at least three times, and the data were analysed using One-Way ANOVA with GraphPad (version 10.1.2).

### Cell culture and viability assay

The HEK293T and HeLa cells were cultured in complete DMEM (Dulbecco’s Modified Eagle’s Medium) high glucose media (Sangon) with 10% FBS (Fetal Bovine Serum, Gibco) and 1% PS (Penicillin-Streptomycin, Sangon) at 37℃ with 5% CO_2_. To test the viability, the cells were seeded in 96-well plate at 10,000/well. After 24 h, the cells were transfected with different concentrations of PNAs as indicated with Lipofectamine 3000 (Thermo Fisher) per the manual. After 72 hours, CCK8 (Sangon) was added to the cells and incubated for 1 h. Then the cell viability is determined by OD 450 measured by the microplate reader (Bioteck).

### qPCR assay

The cells were seeded in 24-well plate at 50,000/well. After 24 h, the cells were transfected with different concentrations of PNA as indicated by Lipofectamine 3000 (Thermo Fisher) per the manual. After 72 h, the total RNAs were extracted by FreeZol Reagent (Vazyme, R711) and reverse transcribed into cDNA using HiScript® III All-in-one RT SuperMix Perfect for qPCR (Vazyme, R333) for pri-miR-21, pre-miR-21 and PTEN mRNA, and miRNA-21 using1st Strand cDNA Synthesis Kit (Vazyme, MR101). qPCR experiment was then conducted using Taq Pro Universal SYBR qPCR Master Mix (Vazyme, Q712) and miRNA Universal SYBR qPCR Master Mix (Vazyme, MQ101) for pre-miR21, pri-miR-21, PTEN mRNA and miRNA, respectively. The experiments were performed on QuantStudio 3 (Applied Biosystems) using the comparative Ct (2^-ΔΔCt^) method. The primers used were synthesized and purified by HPLC (Sangon). DaPNA-198-AAA (NH_2_-Lys-LLTL2TLLAAA-CONH_2_) was applied as a control^18^. The miR-21 expression level is quantified using stem-loop reverse transcription with primer sequence 5′-GTCGTATCCAGTGCAGGGTCCGAGGTATTCGCACTGGATACGACTCAACA-3′. The primer sequence of mature miR-21 used is 5′-GCGCGTAGCTTATCAGACTGA-3′ and 5′-AGTGCAGGGTCCGAGGTATT-3′ and the internal control used is U6 RNA transcript (**Table S2**).

### Western Blot

The cells were seeded in 24-well plate at 50,000/well. After 24 h, the cells were treated as indicated by Lipofectamine 3000 (Thermo Fisher) per the manual. DaPNA-373 (NH_2_-Lys-CAAAAALLLTTL-CONH_2_) was applied as a control. After 72 h, the cell media was removed, washed with 1xPBS (Phosphate Buffer Saline) and the protein was isolated by RIPA with 1% PI (protease inhibitor, Sangon) per the manual. The supernatant containing proteins was collected and determined by BCA Protein Assay Kit (Sangon). 20 µg of protein was loaded on a 10% polyacrylamide gel for SDS-PAGE. The protein was transferred to a 0.4 μM PVDF membrane (Millipore). The membrane was blocked using 5% skim milk (Sangon). Primary antibodies (PTEN and GAPDH, Abcam) were used for incubation with different dilutions per the manual overnight at 4℃ on a shaker. The membrane was washed thrice for a span of 15 mins using 1X TBST (Tris Buffer Saline Tween-20). After being incubated with HRP conjugated secondary antibodies at a dilution of 1:5000 for 1 hour, the membrane was washed thrice before imaging. Efficient chemiluminescent (ECL, Cytiva) solution was used for imaging by Amersham Imager 680. ImageJ software was used for the quantification of the blots.

## ASSOCIATED CONTENT

Supporting Information. HPLC and MS data of PNA oligomers, nondenaturing PAGE and BLI assay data of PNA binding, Dicer cleavage assay data, dual-luciferase reporter assay data.

## AUTHOR INFORMATION

### Author Contributions

Gang Chen, Rongguang Lu, Jean-Luc Decout, Souvik Maiti, and Hongwei Wang conceived this project. Anni Wang, Ruiyu Lin, Xuan Zhan, Alan Ann Lerk Ong, Desiree-Faye Kaixin Toh, Kiran M. Patil, Manchugondanahalli S. Krishna, and Clément Dezanet synthesized the PNA oligomers. Yun Lian performed the PAGE, BLI, dual-luciferase, CCK8, qPCR and Western Blot experiments. Arpita Ghosha and Vignesh Rathinavelpandian Anandavela performed initial cell culture studies. Xin Ke performed the Dicer enzyme cleavage activity assay.

### Notes

A China patent (ZL 2023 1 1137198.3) has been granted based on some of the work reported here. A Patent Cooperation Treaty (PCT) patent application based on some of the work reported here has been filed.

