## Supplemental Information for "A new dual-affinity peptide nucleic acid for targeting miRNA-21 precursor rescues tumor repressor PTEN expression"

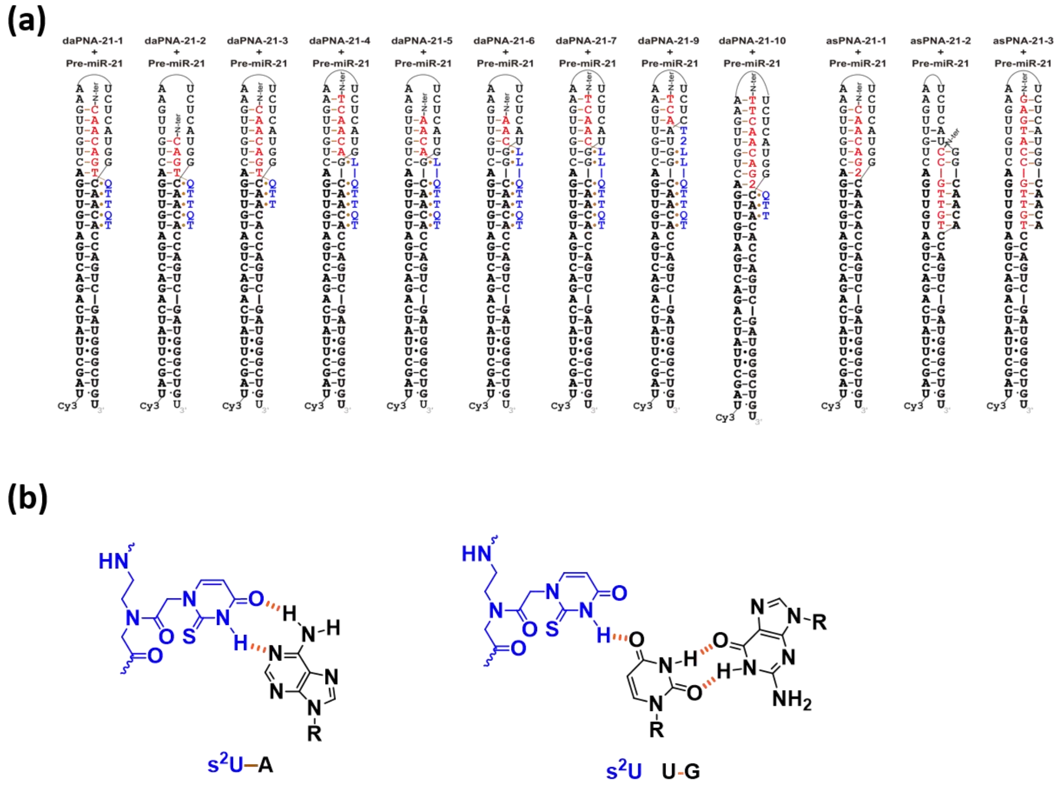


**Figure S1.** Schemes of potential PNA-RNA complex structures. Only daPNA-21-1, daPNA-21-3, daPNA-21-7, and daPNA-21-10 show observable binding (Figure 4c,S3 and Table S1). (a) Schemes of potential PNA-RNA structures for daPNA-1-10, asPNA-21-1-3. The mature miR-21 sequences are shown in bold. (b) Chemical structure of $s^2U$-A Watson-Crick pair and $s^2U{\cdot}U$-G triple.

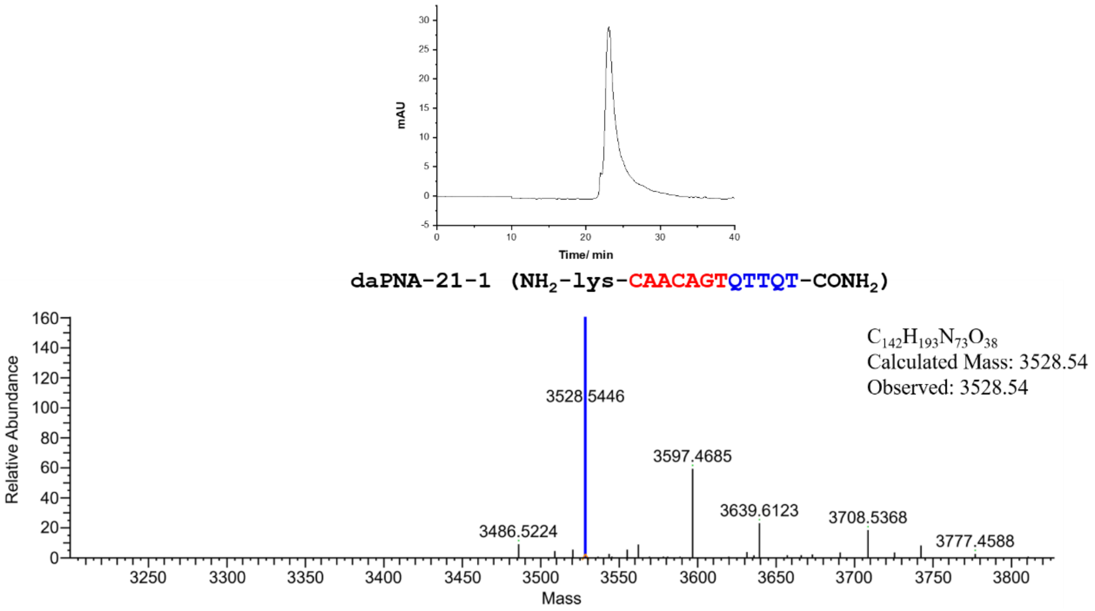


**Figure S2a.** HPLC (top) and LC/MS (bottom) data of daPNA-21-1.

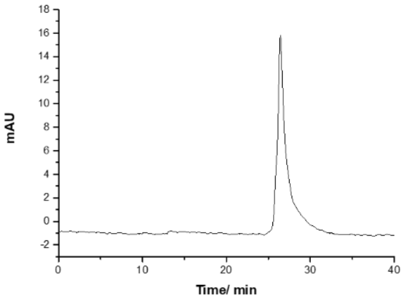


**daPNA-21-2 (NH$_2$-lys-CAGTQTTQT-CONH$_2$)**

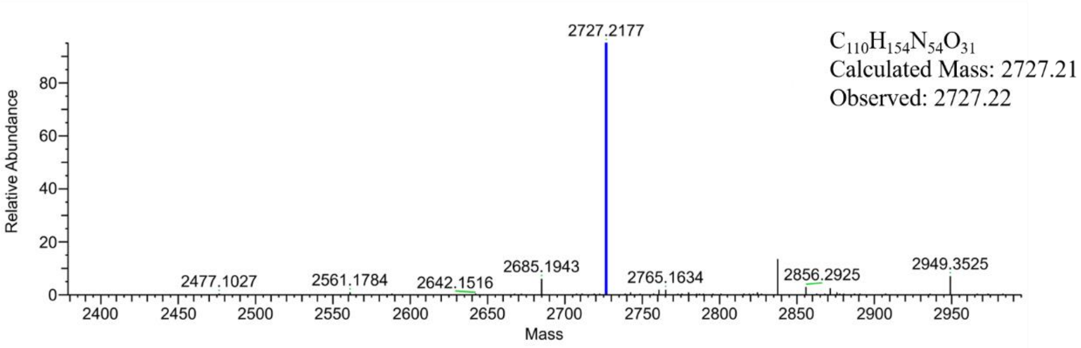


**Figure S2b.** HPLC (top) and LC/MS (bottom) data of daPNA-21-2.

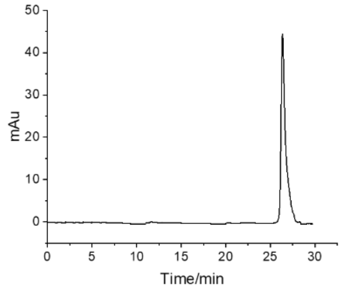


**daPNA-21-3 (NH$_2$-lys-CAACAGTQTT-CONH$_2$)**

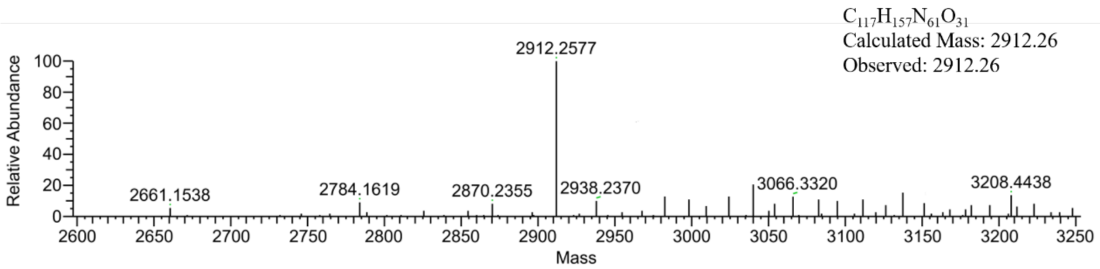


**Figure S2c.** HPLC (top) and LC/MS (bottom) data of daPNA-21-3.

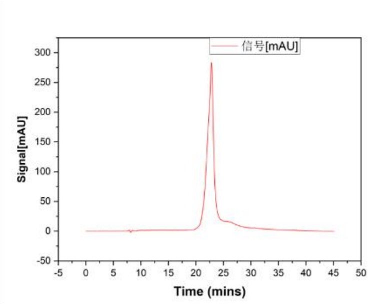


**daPNA-21-4 (NH$_2$-lys-TCAACALQTTQT-CONH$_2$)**

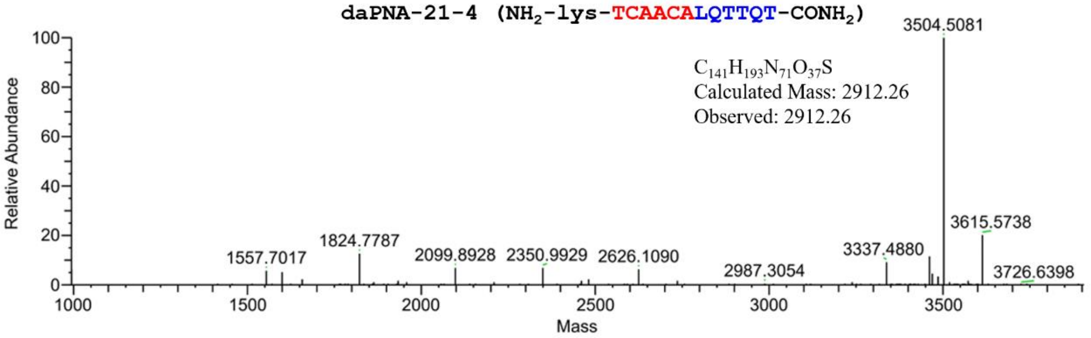


**Figure S2d.** HPLC (top) and LC/MS (bottom) data of daPNA-21-4.

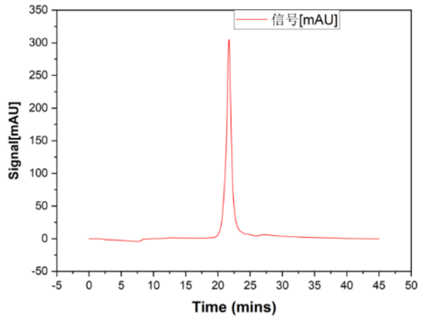


**daPNA-21-5 (NH$_2$-lys-AACALQTTQT-CONH$_2$)**

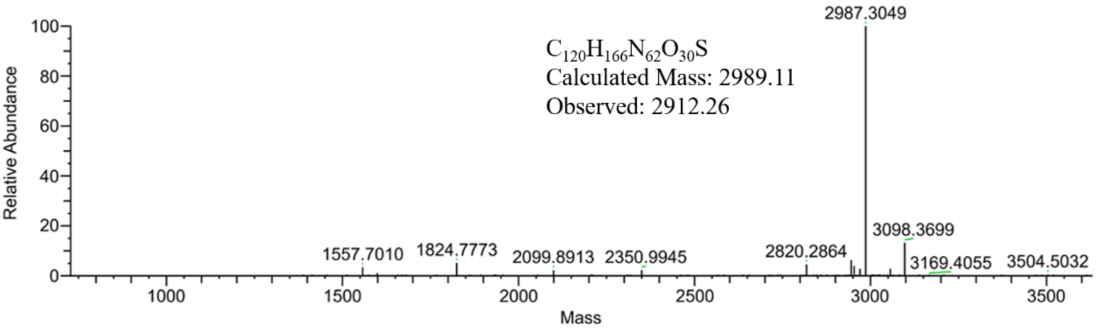


**Figure S2e.** HPLC (top) and LC/MS (bottom) data of daPNA-21-5.

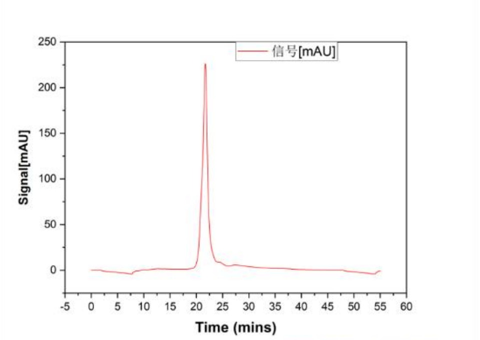


**daPNA-21-6 ($NH_2$-lys-AACLLQTTQT-$CONH_2$)**

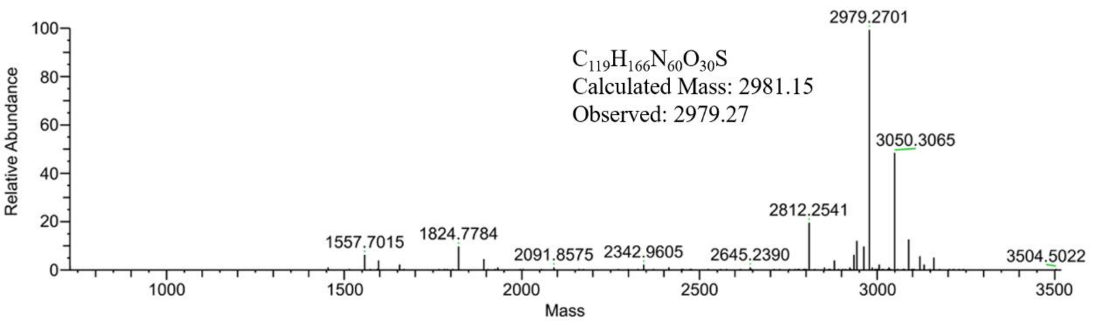


**Figure S2f.** HPLC (top) and LC/MS (bottom) data of daPNA-21-6.

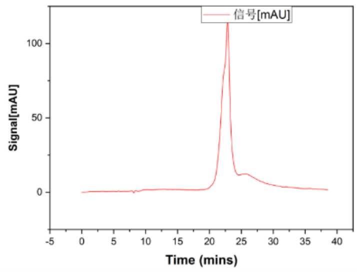


**daPNA-21-7 ($NH_2$-lys-TCAACLLQTTQT-$CONH_2$)**

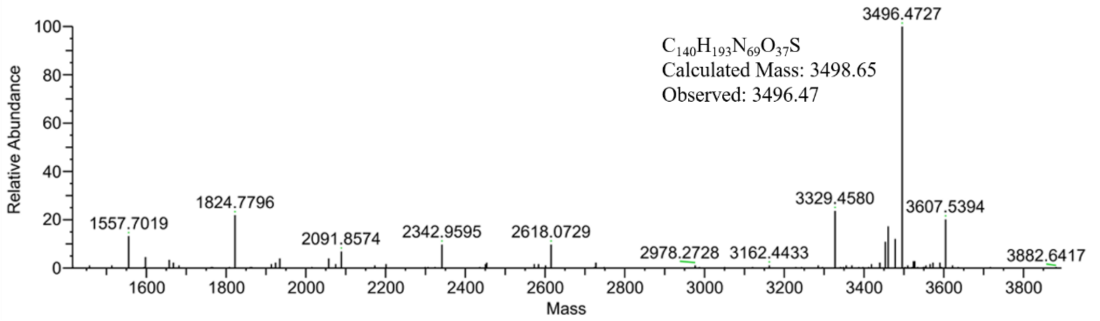


**Figure S2g.** HPLC (top) and LC/MS (bottom) data of daPNA-21-7.

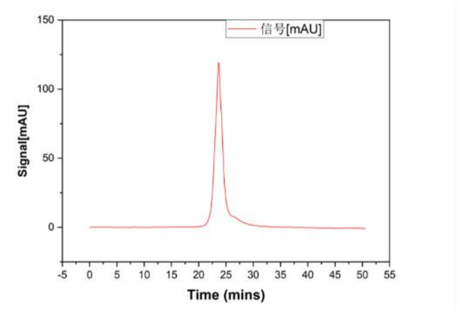


**daPNA-21-9 ($NH_2$-lys-TCAT2LLQTTQT-$CONH_2$)**

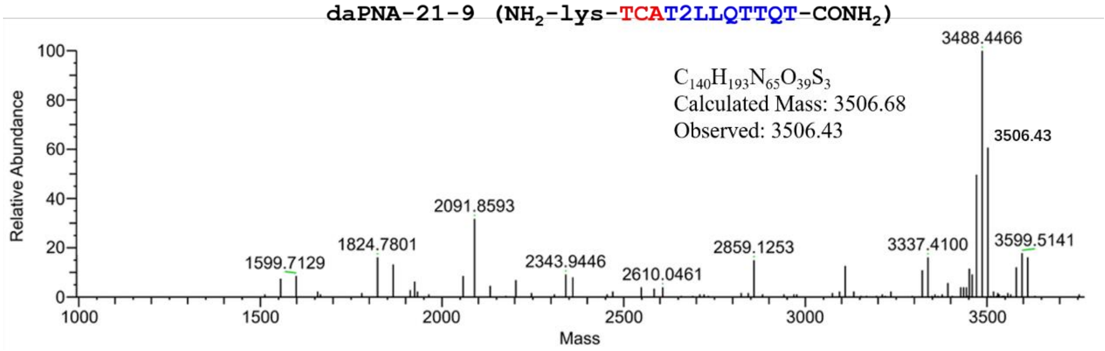


**Figure S2h.** HPLC (top) and LC/MS (bottom) data of daPNA-21-9.

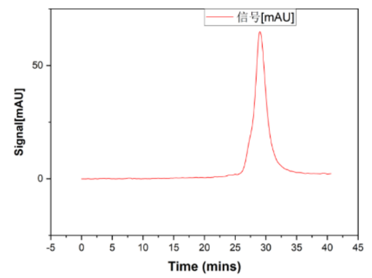


**daPNA-21-10 ($NH_2$-lys-TTCAACAG2QTT-$CONH_2$)**

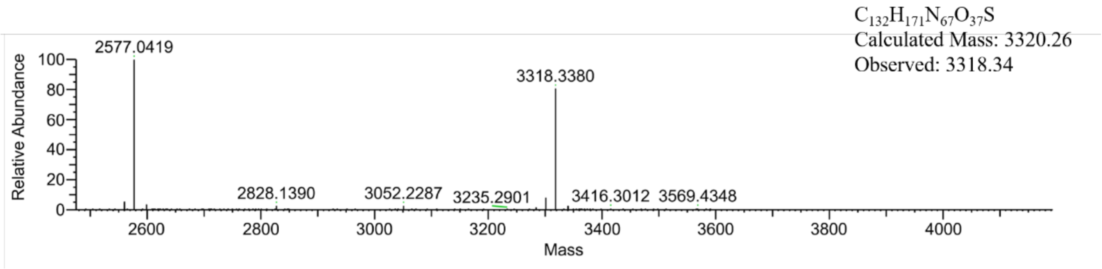


**Figure S2i.** HPLC (top) and LC/MS (bottom) data of daPNA-21-10.

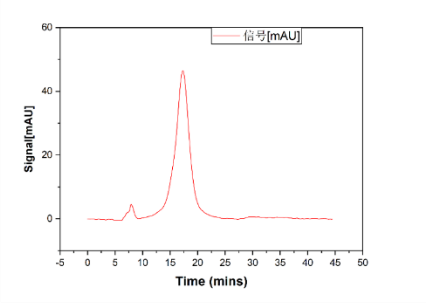


**asPNA-21-1 (NH$_2$-lys-CAACAG2-CONH$_2$)**

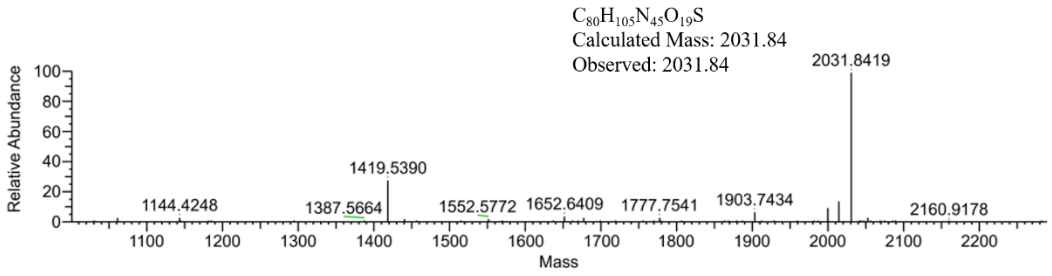


**Figure S2j.** HPLC (top) and LC/MS (bottom) data of asPNA-21-1.

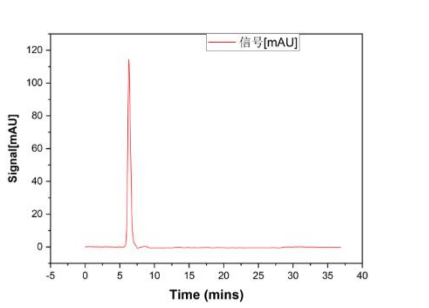


**asPNA-21-2 (NH$_2$-lys-CCGTTGT-CONH$_2$)**

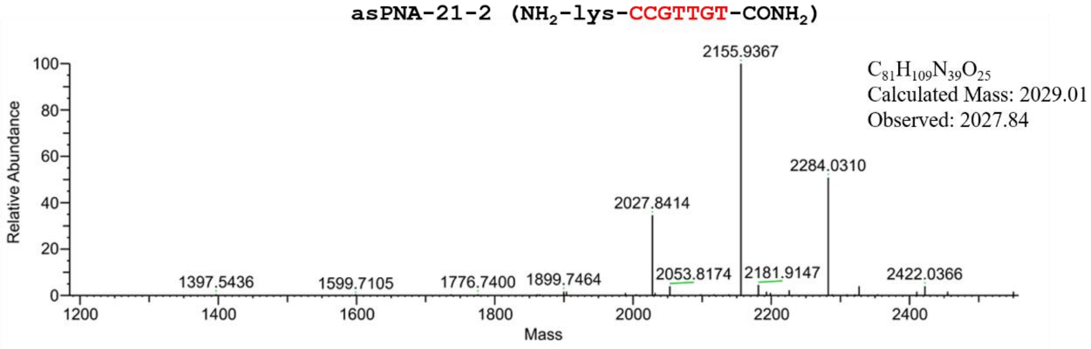


**Figure S2k.** HPLC (top) and LC/MS (bottom) data of asPNA-21-2.

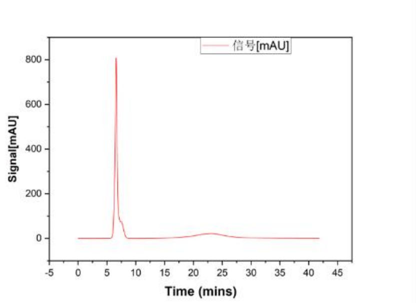


**asPNA-21-3 ($NH_2$-lys-GAGTACCGTTGT-$CONH_2$)**

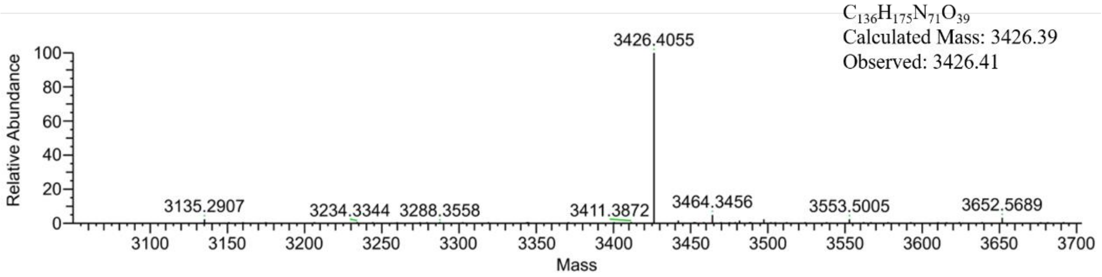


**Figure S2l.** HPLC (top) and LC/MS (bottom) data of asPNA-21-3.

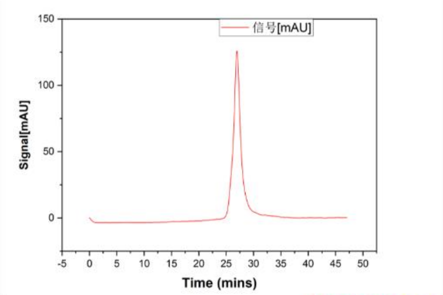


**daPNA-373-1 ($NH_2$-lys-CAAAAALLLTTL-$CONH_2$)**

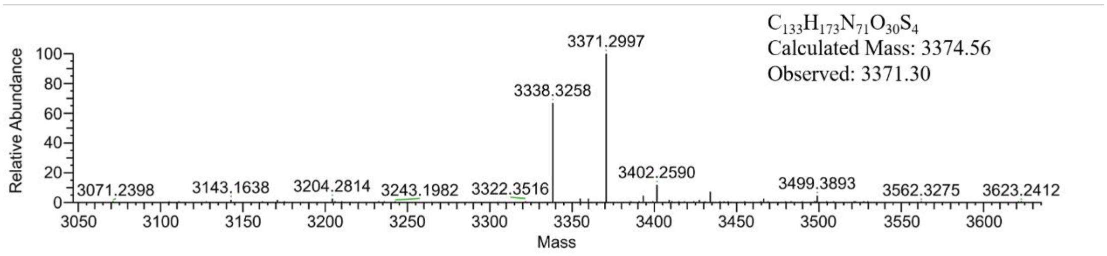


**Figure S2m.** HPLC (top) and LC/MS (bottom) data of daPNA-373-1.

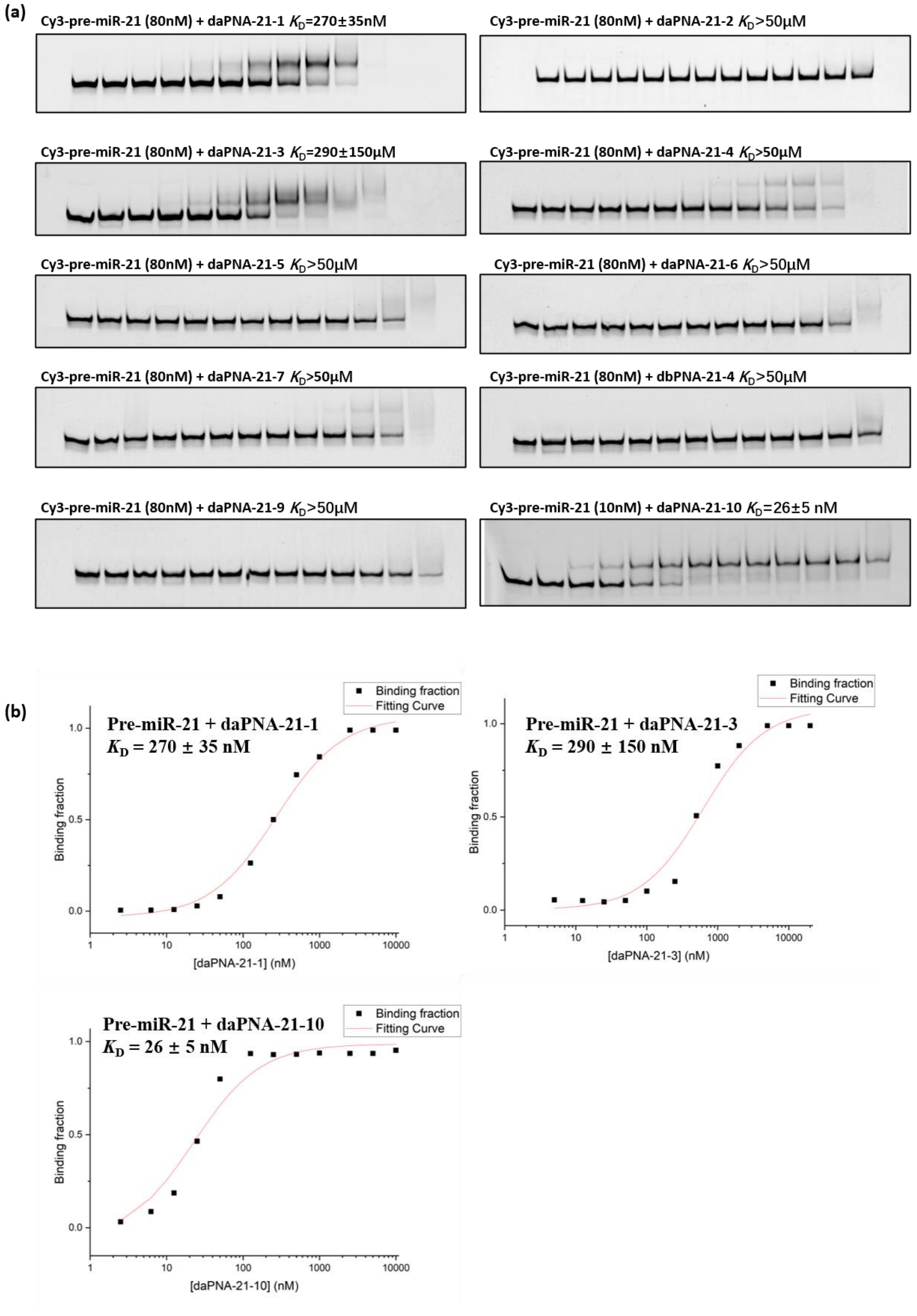


**Figure S3.** PAGE data for the quantification of binding affinities of pre-miR-21 with daPNAs. The final concentrations of daPNAs loaded are 0, 2, 6, 12, 25, 50, 125, 250, 500, 1,000, 2,500, 5,000, and 10,000 nM, respectively. (a) PAGE gel for daPNA-21s. (b) Fitting results for daPNA-21-1, 3, 10.

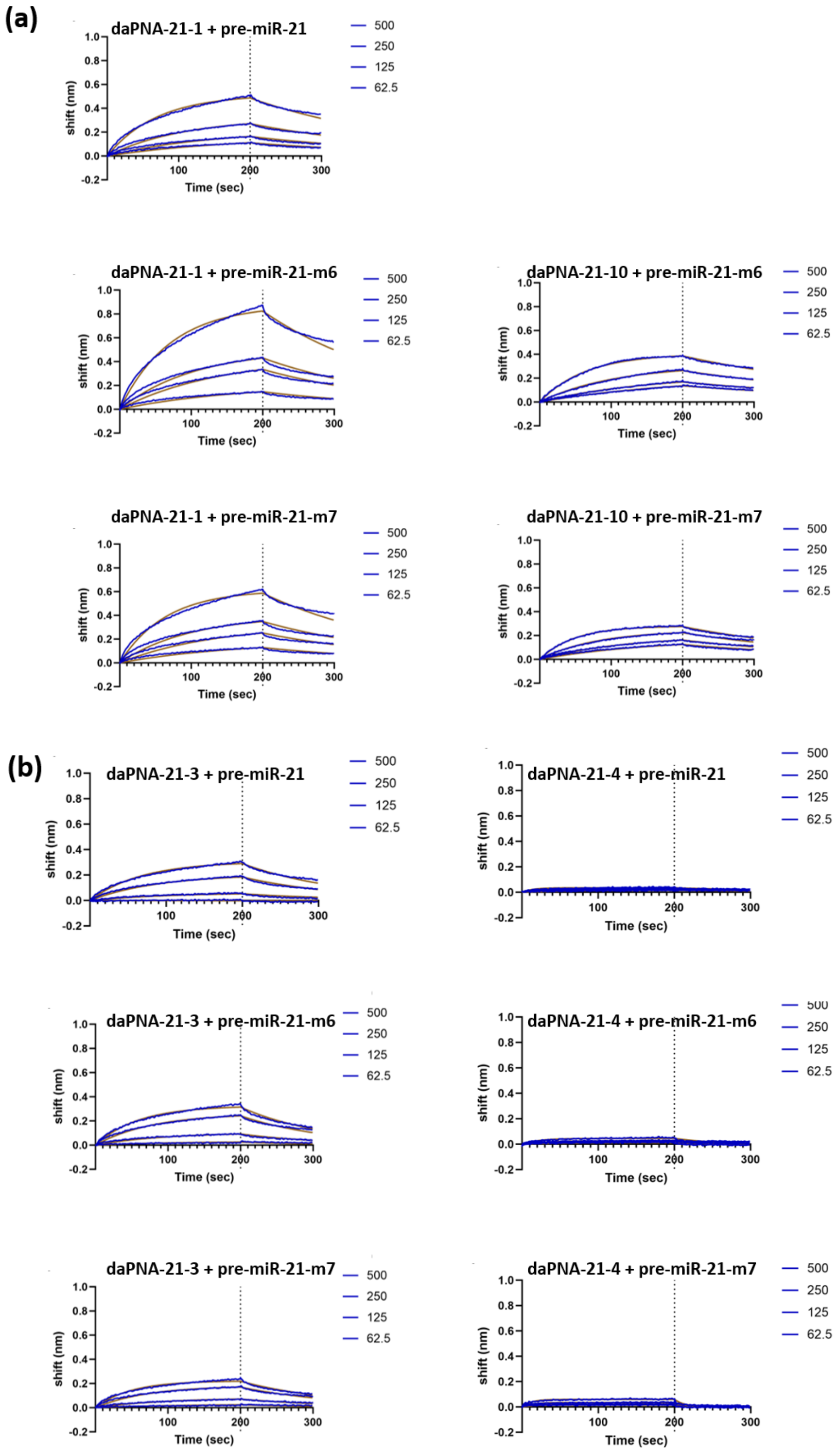
(a)
daPNA-21-1 + pre-miR-21
daPNA-21-1 + pre-miR-21-m6
daPNA-21-10 + pre-miR-21-m6
daPNA-21-1 + pre-miR-21-m7
daPNA-21-10 + pre-miR-21-m7
(b)
daPNA-21-3 + pre-miR-21
daPNA-21-4 + pre-miR-21
daPNA-21-3 + pre-miR-21-m6
daPNA-21-4 + pre-miR-21-m6
daPNA-21-3 + pre-miR-21-m7
daPNA-21-4 + pre-miR-21-m7
500
250
125
62.5
shift (nm)
Time (sec)
1.0
0.8
0.6
0.4
0.2
0.0
-0.2
100
200
300

(c)

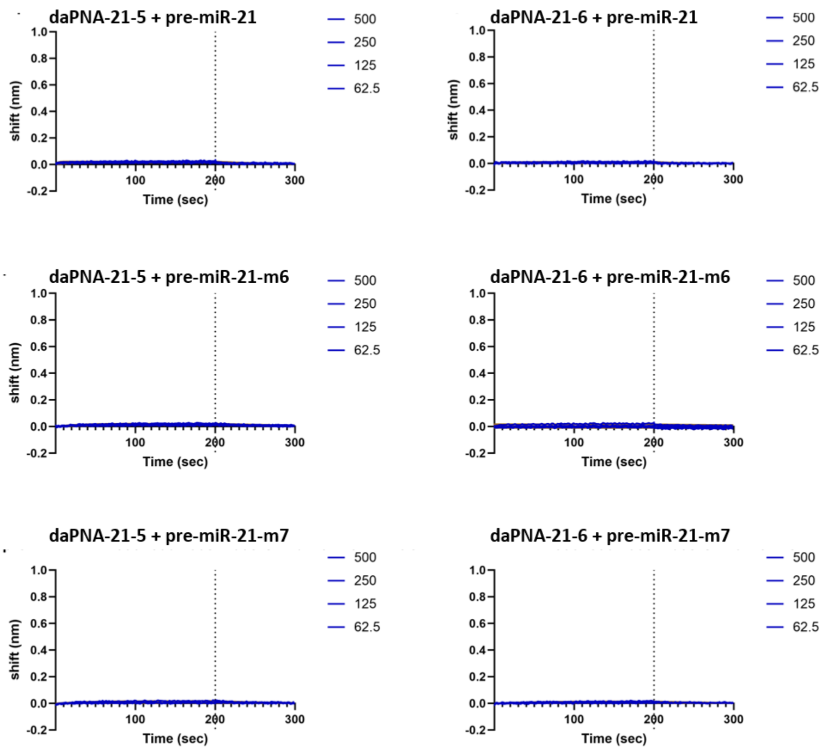
daPNA-21-5 + pre-miR-21
daPNA-21-6 + pre-miR-21
daPNA-21-5 + pre-miR-21-m6
daPNA-21-6 + pre-miR-21-m6
daPNA-21-5 + pre-miR-21-m7
daPNA-21-6 + pre-miR-21-m7
shift (nm)
Time (sec)
500
250
125
62.5

(d)

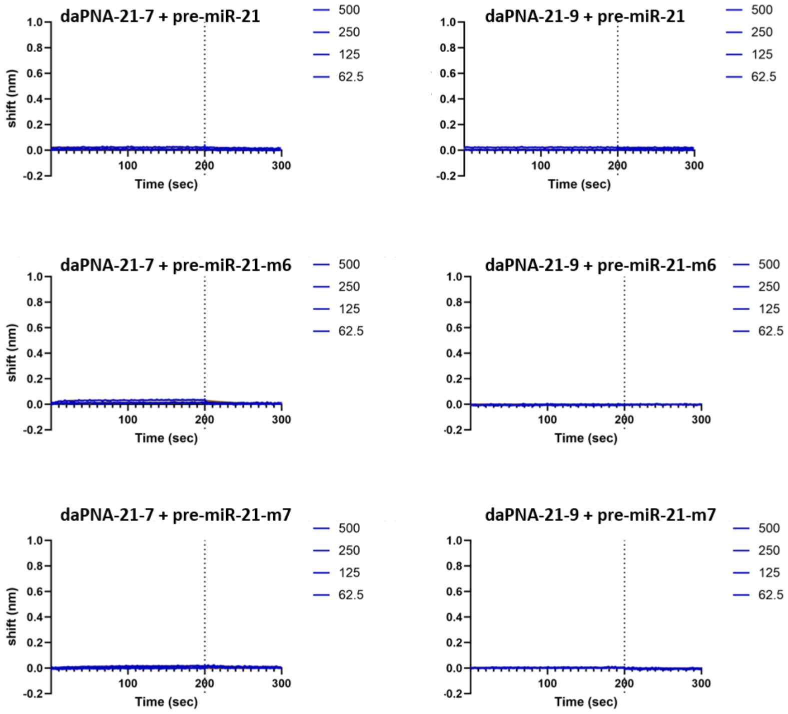
daPNA-21-7 + pre-miR-21
daPNA-21-9 + pre-miR-21
daPNA-21-7 + pre-miR-21-m6
daPNA-21-9 + pre-miR-21-m6
daPNA-21-7 + pre-miR-21-m7
daPNA-21-9 + pre-miR-21-m7
shift (nm)
Time (sec)
500
250
125
62.5

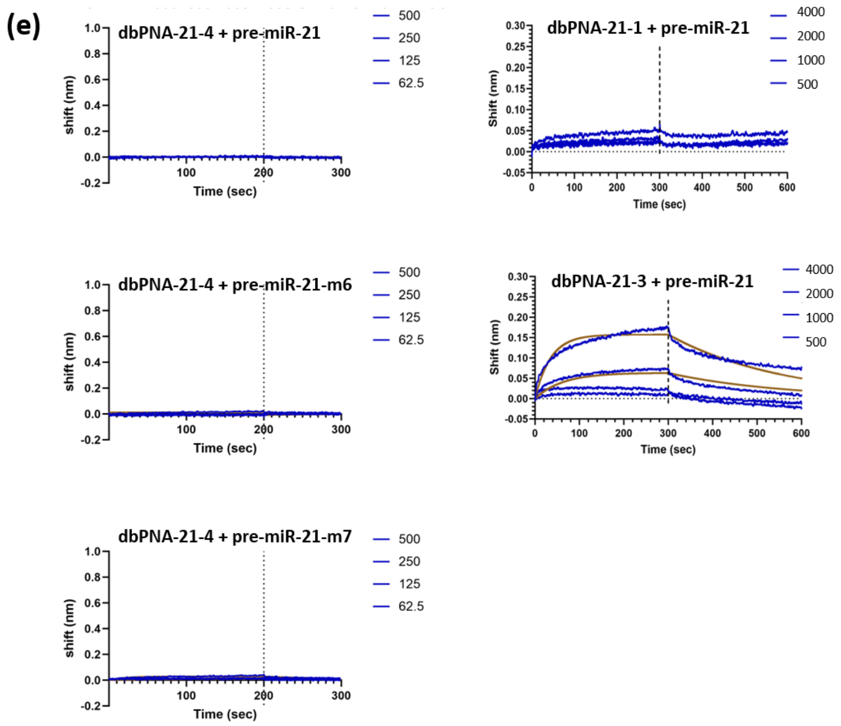


**Figure S4.** BLI data for PNAs binding to biotin-pre-miR-21 and mutants.

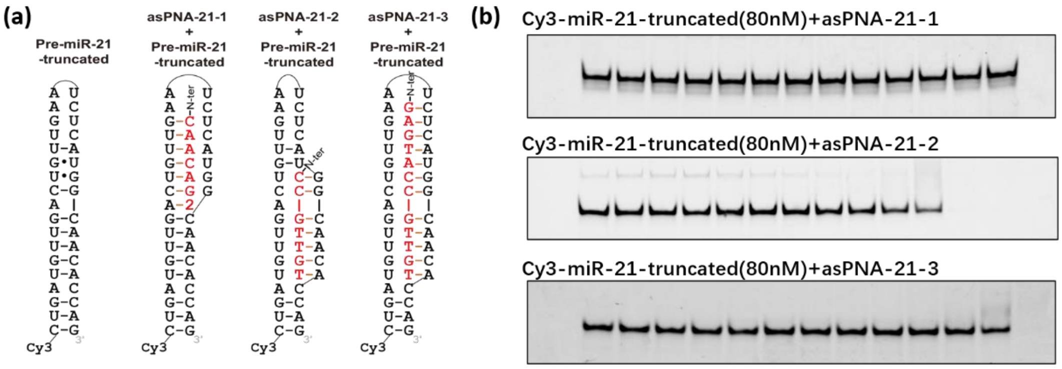


**Figure S5.** PAGE data for the binding of pre-miR-21-truncated with an asPNA. (a) Scheme of asPNA-RNA-truncated complex. (b) PAGE for asPNA-21-1,2,3's titration to Cy3-miR-21-truncated. The final concentrations of asPNAs loaded are 0, 5, 10, 25, 50, 100, 250, 500, 1,000, 2,500, 5,000, 10,000 and 20,000 nM, respectively.

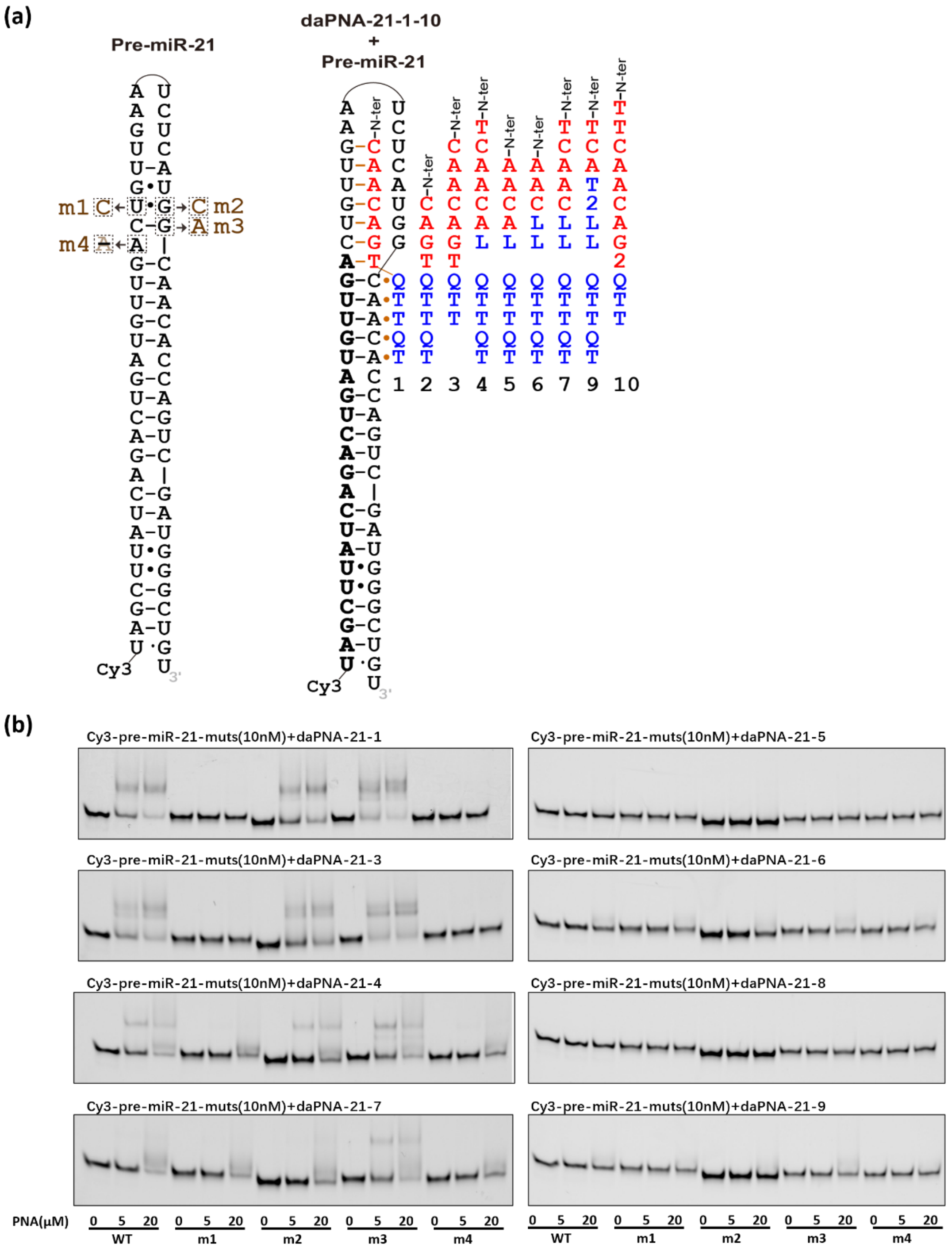


**Figure S6.** PAGE for binding of pre-miR-21-muts with daPNA21-1, 3-9.

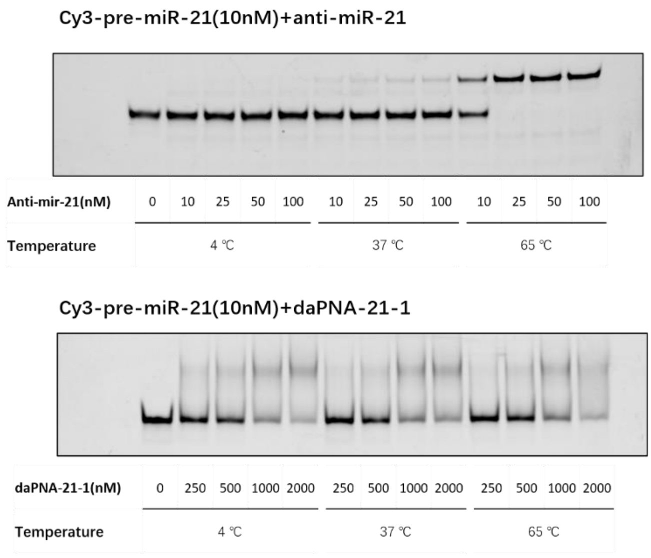


**Figure S7.** Influence of incubation temperature on PNA binding affinity. The PAGE was run at 4 ℃.

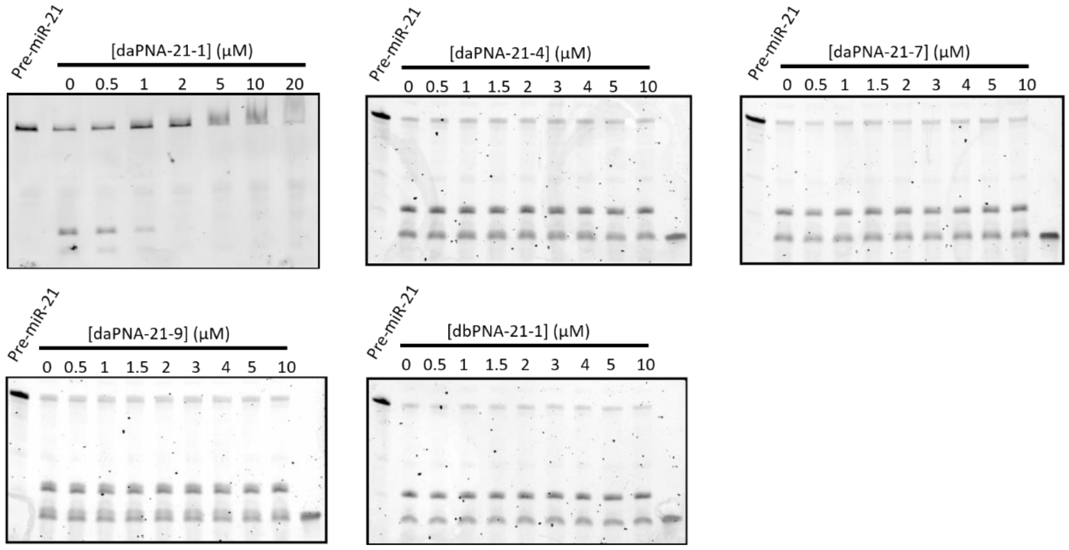


**Figure S8.** Dicer cleavage assay data for pre-miR-21 and effects of daPNA-21-4, daPNA-21-7, daPNA-21-9 and dbPNA-21-1.

(a)

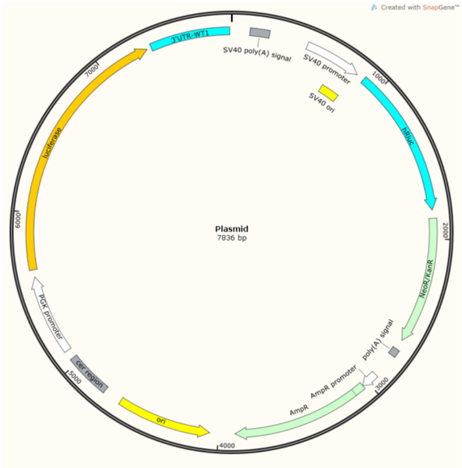


(b)

```
5′...UAUUGAUUGAUUGCUCAUAAGUUAAAUUG...3′      PTEN 3′UTR
          ||::  |    ||  |||||:||
3′-AGUUGUAGUCA--GACUAUUCGAU-5′               miR-21
```

5′...UAGAG**UCAACAUCAGUCUGAUAAGCUA**AAUUG...3′ Plasmid

```
          ||||||||||||||||||||||
      3′-AGUUGUAGUCAGACUAUUCGAU-5′          miR-21
```

(c)

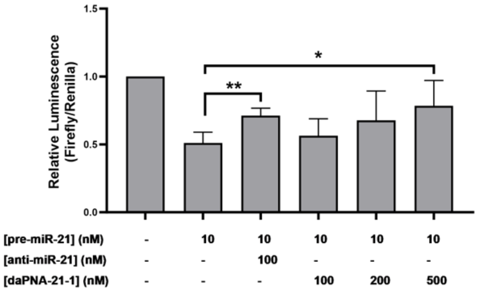


**Figure S9.** Dual-luciferase assay of daPNA-21-1 in HEK293T cells. (a) Scheme of plasmid for dual-luciferase assay. (b) Sequence of binding sites in PTEN mRNA 3′ UTR and plasmid for miR-21. (c) Results of relative luminescence in HEK293T. Cells were transfected with different concentrations of daPNA-21-1 as indicated with pre-miR-21 at 10 nM and treated for 48 h in 96-well plate. The final concentration of the transfected plasmid is 1ng/μl. The variance is analyzed using One-Way ANOVA. $P<0.05$ is considering significant.

**Table S1.** BLI data for daPNAs and asPNAs targeting biotin-pre-miR-21 and mutated hairpin constructs.

| PNA | RNA | $K_D$ (nM) | $k_{on}$ ($M^{-1}$ $s^{-1}$) | $k_{off}$ ($\times 10^{-6}$ $s^{-1}$) | $R^2$ |
|---|---|---|---|---|---|
| daPNA-21-1 | pre-miR-21 | 147±3 | 46,653±921 | 6,877±71 | 0.96 |
| daPNA-21-1 | pre-miR-21-m6 | 408±11 | 20,642±503 | 8,429±70 | 0.98 |
| daPNA-21-1 | pre-miR-21-m7 | 444±19 | 34,127±1,394 | 15,150±165 | 0.94 |
| daPNA-21-10 | pre-miR-21 | 25±1 | 388,793±10,254 | 9,564±119 | 0.95 |
| daPNA-21-10 | pre-miR-21-m6 | 177±2 | 32,686±348 | 5,785±38 | 0.99 |
| daPNA-21-10 | pre-miR-21-m7 | 120±1 | 35,603±282 | 4,276±30 | 0.99 |
| daPNA-21-3 | pre-miR-21 | 535±13 | 14,242±322 | 7,622±51 | 0.99 |
| daPNA-21-3 | pre-miR-21-m6 | 504±10 | 16,928±328 | 8,535±48 | 0.99 |
| daPNA-21-3 | pre-miR-21-m7 | 275±5 | 26,733±407 | 7,361±50 | 0.99 |
| daPNA-21-4 | pre-miR-21 | No binding | | | |
| daPNA-21-4 | pre-miR-21-m6 | No binding | | | |
| daPNA-21-4 | pre-miR-21-m7 | No binding | | | |
| daPNA-21-5 | pre-miR-21 | No binding | | | |
| daPNA-21-5 | pre-miR-21-m6 | No binding | | | |
| daPNA-21-5 | pre-miR-21-m7 | No binding | | | |
| daPNA-21-6 | pre-miR-21 | No binding | | | |
| daPNA-21-6 | pre-miR-21-m6 | No binding | | | |
| daPNA-21-6 | pre-miR-21-m7 | No binding | | | |
| daPNA-21-7 | pre-miR-21 | No binding | | | |
| daPNA-21-7 | pre-miR-21-m6 | No binding | | | |
| daPNA-21-7 | pre-miR-21-m7 | No binding | | | |
| daPNA-21-8 | pre-miR-21 | No binding | | | |
| daPNA-21-8 | pre-miR-21-m6 | No binding | | | |
| daPNA-21-8 | pre-miR-21-m7 | No binding | | | |
| daPNA-21-9 | pre-miR-21 | No binding | | | |
| daPNA-21-9 | pre-miR-21-m6 | No binding | | | |
| daPNA-21-9 | pre-miR-21-m7 | No binding | | | |
| dbPNA-21-1 | pre-miR-21 | No binding | | | |
| dbPNA-21-1 | pre-miR-21-m6 | No binding | | | |
| dbPNA-21-1 | pre-miR-21-m7 | No binding | | | |
| asPNA-21-1 | pre-miR-21 | No binding | | | |
| asPNA-21-1 | pre-miR-21-m6 | No binding | | | |
| asPNA-21-1 | pre-miR-21-m7 | No binding | | | |
| dbPNA-21-4 | pre-miR-21 | No binding | | | |
| dbPNA-21-4 | pre-miR-21-m6 | No binding | | | |
| dbPNA-21-4 | pre-miR-21-m7 | No binding | | | |
| dbPNA-21-1 | pre-miR-21 | No binding | | | |
| dbPNA-21-3 | pre-miR-21 | 623±9 | 6,160±23 | 3,840±20 | 0.99 |

**Table S2.** Primers used for qPCR.

| Primer | Sequence |
|---|---|
| Hsa-GAPDH-F | GGAGCGAGATCCCTCCAAAAT |
| Hsa-GAPDH-R | GGCTGTTGTCATACTTCTCATGG |
| Hsa-PTEN-F | TTTGAAGACCATAACCCACCAC |
| Hsa-PTEN-R | ATTACACCAGTTCGTCCCTTTC |
| Hsa-pre-miR-21-F | CTGATGTTGACTGTTGA |
| Hsa-pre-miR-21-R | GCCCATCGACTGGTGTTGCC |
| miR21-F | GCGCGTAGCTTATCAGACTGA |
| miR21-R | AGTGCAGGGTCCGAGGTATT |
| U6-F | CTCGCTTCGGCAGCACA |
| U6-R | AACGCTTCACGAATTTGCGT |
| Stemloop-21Rev GTCGTATCCAGTGCAGGGTCCGAGGTATTCGCACTGGATACGACTCAACA | |
| miR373-F | CGACTCAAAATGGGGGCG |
| miR373-R | AGTGCAGGGTCCGAGGTATT |
| Stemloop-373Rev GTCGTATCCAGTGCAGGGTCCGAGGTATTCGCACTGGATACGACGGAAAG | |
